# Hemozoin induces Malaria via activation of DNA damage, p38 MAPK and Neurodegenerative Pathways in Human iPSC-derived Neuronal Model of Cerebral Malaria

**DOI:** 10.1101/2024.03.08.584068

**Authors:** Abida lslam Pranty, Leon-Phillip Szepanowski, Wasco Wruck, Akua Afriyie Karikari, James Adjaye

## Abstract

Malaria caused by *Plasmodium falciparum* infection results in severe complications including cerebral malaria (CM), in which approximately 30% of patients end up with neurological sequelae. Sparse in vitro cell culture-based experimental models which recapitulate the molecular basis of CM in humans has impeded progress in our understanding of its etiology. This study employed healthy human induced pluripotent stem cells (iPSCs) derived neuronal cultures stimulated with hemozoin (HMZ)-the malarial toxin as a model for CM. Secretome, qRT-PCR, Metascape, and KEGG pathway analyses were conducted to assess elevated proteins, genes, and pathways. Neuronal cultures treated with HMZ showed enhanced secretion of interferon-gamma (IFN-γ), interleukin (IL)1-beta (IL-1β), IL-8 and IL-16. Enrichment analysis revealed malaria, positive regulation of cytokine production and positive regulation of mitogen-activated protein kinase (MAPK) cascade which confirm inflammatory response to HMZ exposure. KEGG assessment revealed up-regulation of malaria, MAPK and neurodegenerative diseases-associated pathways which corroborates findings from previous studies. Additionally, HMZ induced DNA damage in neurons. This study has unveiled that exposure of neuronal cultures to HMZ, activates molecules and pathways similar to that observed in CM and neurodegenerative diseases. Furthermore, our model is an alternative to rodent experimental models of CM.

## Introduction

In spite of the significant efforts to combat malaria, the disease remains a public health problem globally. In 2021, there were an estimated 247 million malaria cases in 84 malaria endemic countries, which represents an increase of 2 million cases compared with 2020^1^. Also, the continuous importation of malaria to non-endemic regions consistently challenges elimination efforts, leading to the spread of drug-resistant parasites within and between countries^2^. *Plasmodium falciparum* and *vivax* are the most virulent of the five human parasite species. Nonetheless, *P. falciparum* is the species primarily implicated in severe and fatal forms of clinical malaria^3^. Principal to the pathogenesis of *P. falciparum* is the toxin, hemozoin (HMZ), which is also referred to as the malaria pigment. Falciparum does not infiltrate the brain parenchyma but can cause severe damage to neurons, through the release of HMZ, which can induce reactive oxygen species (ROS) mediated macromolecular damage and inflammatory response^4,5^. Moreover, *P. falciparum* poses a challenge for experimental malaria investigations, due to its inability to infect species like rodents^6^. A majority of falciparum malaria research is based on clinical studies, which limits access to certain tissues such as the brain. Human derived induced pluripotent stem cells (iPSCs) have emerged as a relevant tool for modelling several diseases *in-vitro*^7,8^. iPSCs can be used for extensive studies related to disease pathogenesis, the impact of exogenous factors and the molecular targets of disease-modulating treatments. With the lack of an appropriate experimental model for *P. falciparum*, and with the difficulty in studying the molecular-basis and mechanisms associated with malaria in human subjects, there is a pressing need for new models of falciparum malaria.

A major complication of falciparum infection is cerebral malaria (CM). Clinically, CM is characterized by lack of consciousness and coma, with high mortality rates^9^. Post-CM survivors sustain brain injuries and neurological sequelae including cognitive impairment, motor skill impairment, cortical blindness, seizures and attention deficit hyperactive disorder^9,10^. To date, the pathogenic mechanisms leading to cerebral malaria remains an enigma as studies have been hampered by limited accessibility to human tissues and brain injury in human post-mortem tissues has provided limited cross-sectional data^11^. In a previous study using pooled transcriptome data from whole blood analysis of *P. falciparum* malaria patients, we discovered that falciparum malaria shared similar biological processes with neurodegenerative diseases^4^. Additionally, in our previous study we employed an iPSC-based model discovering that malaria was a significant pathway in constructing a network of genes associated with late onset Alzheimer’s disease (LOAD)^12^. Both mild and severe forms of falciparum malaria have been previously linked to psychiatric disorders such as depression, irritability, anxiety, difficulties with concentration, disorientation and forgetfulness^13^. Deciphering the molecular changes that occur during the disease pathogenesis could unravel the link between falciparum malaria and neurodegenerative diseases.

In this study, we employed two healthy iPSC-line-derived neuronal cultures to characterize molecular changes that occur upon exposure of neuronal cultures to the parasite’s most potent toxin-HMZ. After exposure to HMZ, qRT-PCR and secretome analysis revealed an increase in pro-inflammatory molecules and neurotrophins including interleukin-1-beta (IL-1β), IL-16, interferon-gamma (IFN-γ), monocyte chemotactic protein-1 (MCP-1), vascular endothelial growth factor (VEGF) and brain derived neurotrophic factor (BDNF). Pathway enrichment analysis using the Kyoto Encyclopedia of Genes and Genomes (KEGG) network revealed malaria, mitogen-activated protein kinase (MAPK), neurodegenerative and Alzheimer’s disease (AD) pathways, as upregulated in HMZ-exposed cells. This corroborates our previous study where neurodegenerative and AD pathways were elevated in falciparum malaria patients. Metascape assessment uncovered malaria, regulation of leukocyte proliferation, positive regulation of cytokine production and positive regulation of MAPK cascade, which are indicative of cellular inflammatory response to HMZ exposure. Furthermore, western blot analysis confirmed DNA damage in UM51-derived neurons, implicating HMZ as genotoxic. Investigations using HMC3 microglia cells revealed a similar pattern of inflammation and cellular injury. Taken together, the findings of this study consolidate the theory that CM and neurodegenerative diseases, such as AD, activate similar cellular mechanisms through inflammation, which could account for the neurological deficits observed in CM patients. It also demonstrates that iPSC-derived neuronal cultures could serve as a valuable model in research regarding CM.

## Results

### Generation of iPSC-derived neuronal networks to model cerebral malaria

The UM51 line was generated from SIX2-positive renal progenitor cells isolated from the urine of a 51-year-old healthy male of African origin^14^, while healthy human fetal foreskin fibroblasts were reprogrammed to generate the B4-iPSC line (HFF1, ATCC, #ATCCSCRC-1041, http://www.atcc.org)^15^ (Supplementary Table S1). To obtain neuronal networks, UM51 and B4 iPSCs were seeded as single cells for neural induction, then expanded as neural progenitor cells and further differentiated into neurons (Fig. 1a, Supplementary Fig. 1). A dose-response curve was prepared using ascending concentrations of HMZ on neuronal cultures to determine IC80 (cell survival rate) and the closest lower concentration examined (20 μM) was used for all following experiments (Supplementary Fig. 1). Subsequently, day 16 neuronal cultures from both cell lines were exposed to 20 μM hemozoin (HMZ) for 48 h to observe HMZ-induced effects in central nervous system (CNS) cell populations with the aim of modeling cerebral malaria using iPSC-derived neuronal cultures (Fig. 1a, Supplementary Fig. 1).

**Figure 1.**
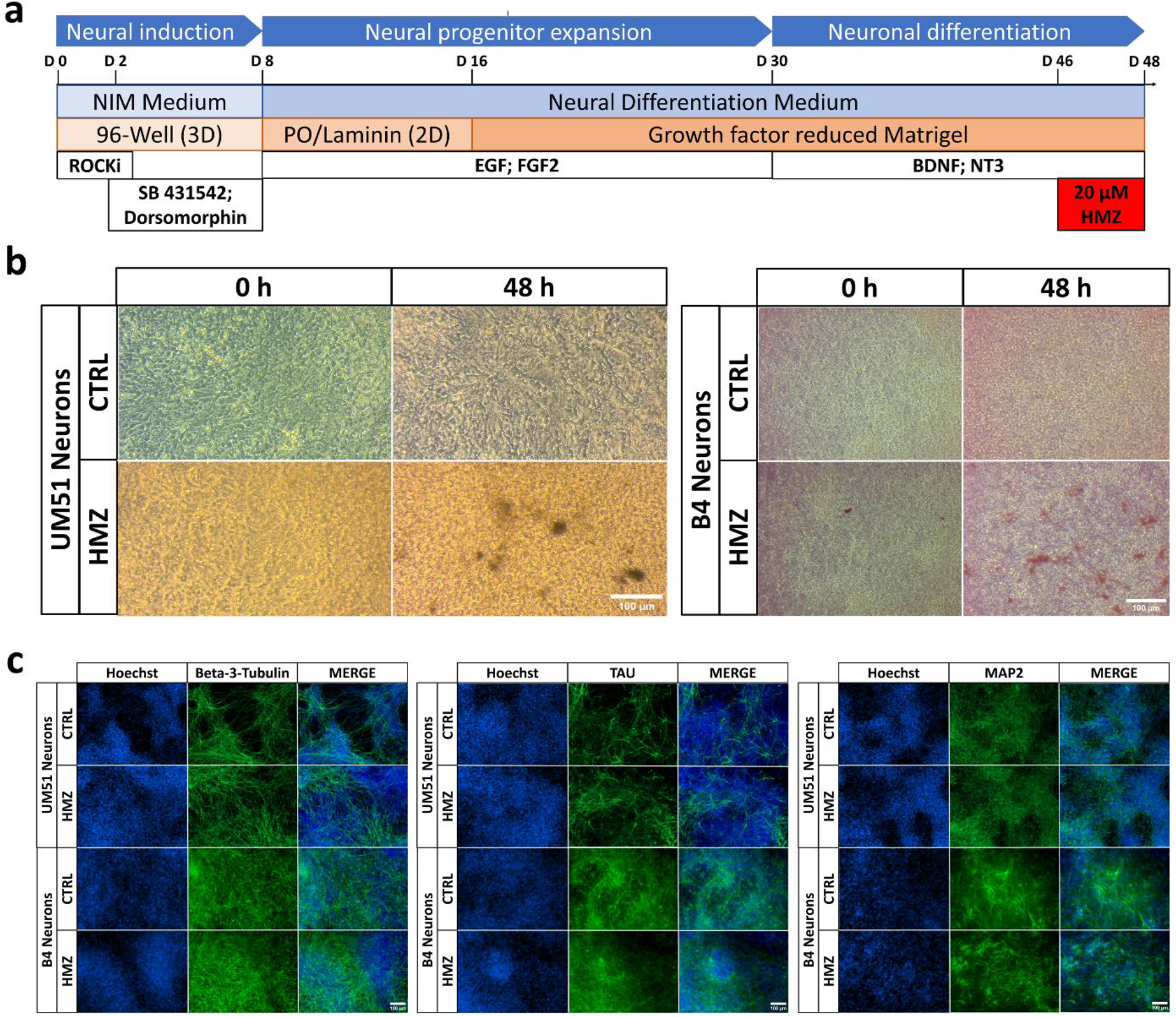
Generation, characterization and utilization of iPSC-derived neuronal cultures for modelling cerebral malaria. **(a)** Schematic outline of the protocol used to generate iPSC-derived neuronal cultures. **(b)** Representative bright field images of control and HMZ-treated UM51- and B4 iPSC-derived neuronal cultures at 0h and 48 h of exposure. Scale bar 100 μm. **(c)** Representative immunocytochemistry (ICC) images of β3-Tubulin-, TAU- and MAP2-positive cells in UM51- and B4-derived neuronal cultures after 48 h HMZ exposure in comparison to control. Scale bar 100 μm.

Bright field microscopy showed deposition of malaria pigments as brown crystals inside or around the neuronal cultures after 48 h of treatment (Fig. 1b, Supplementary Fig. 1). Neuronal features were confirmed in both control and HMZ-treated neuronal cultures by expression of neuronal markers Beta III tubulin (β3-Tubulin), tubulin associated unit (TAU), and microtubule-associated protein 2 (MAP2) (Fig. 1c, Supplementary Fig. 1).

### HMZ activates inflammatory-associated pathways in iPSC-derived neuronal cultures

Supernatants from UM51 neuronal cultures were used to analyze the secretome profile of both HMZ-treated and control conditions (Supplementary Fig. 2). The resulting secretome profiles were further analyzed utilizing hierarchical clustering analysis (heatmap using Pearson correlation as similarity measure in Fig. 2a), which indicated induction of an inflammatory response upon treatment with 20 μM HMZ for 48 h. Usually pro- and anti-inflammatory cytokines are involved not only in inflammatory response but also in immune-regulatory functions. A number of crucial pro-inflammatory cytokines such as IL-8, IL1-B, IFN-G, IL-16 showed enhanced secretion, while anti-inflammatory cytokines such as IL-4, IL-13 showed reduced secretion (Fig. 2a) (Supplementary Fig. 2).

**Figure 2.**
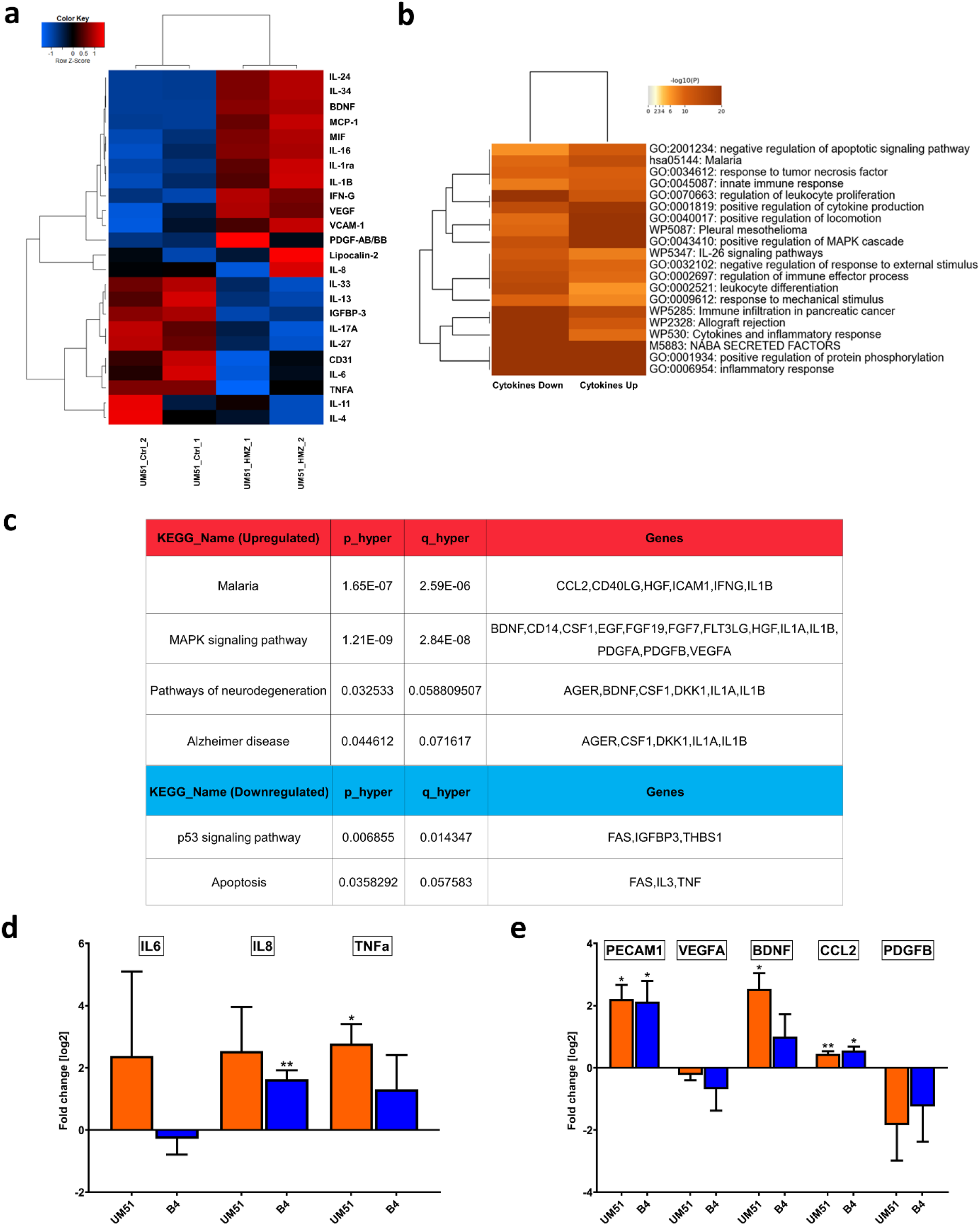
Secretome analysis of neuronal cultures upon HMZ exposure reveals activation of inflammatory response, MAPK-signaling and neurodegenerative pathway. Cytokine array analyses was performed with supernatant derived from the UM51 neuronal cultures. **(a)** Pearson’s heatmap depicting selected chemo- and cytokines regulated in UM51 neuronal cultures after 48 h HMZ exposure in comparison to control. **(b)** Metascape-generated heatmap comparing the sets of up- and downregulated chemo- and cytokines derived from UM51 neuronal cultures after 48 h HMZ exposure reveals a secretome signature involved in i.a. Malaria, response to tumor necrosis factor and positive regulation of MAPK cascade. **(c)** Table depicting selected KEGG pathways including up-regulated malaria, MAPK signalling pathway, pathways of neurodegeneration and Alzheimer disease and down-regulated p53 signalling pathway and apoptosis, as well as associated genes upon HMZ exposure in comparison to control. **(d)** Relative mRNA expression analysis of *IL6, IL8* and *TNFA* in UM51 and B4-derived neuronal cultures after 48 h HMZ exposure in comparison to control. **(e)** Relative mRNA expression analysis of *PECAM1, VEGFA, BDNF, CCL2* and *PDGFB* in UM51 and B4-derived neuronal cultures after 48 h HMZ exposure in comparison to control. (d,e) UM51 n=3; B4 Ctrl n=2; B4 HMZ n=3; blots depict mean and error bars depict SD of all experiments. Asterisk (*) depicts significance, which is indicated by *p<0.05; **p<0.01.

We further performed a Metascape-based analysis utilizing the list of up- and down-regulated cytokines obtained from the UM51 neuronal culture. The resulting enrichment clusters revealed malaria, positive regulation of cytokine production, positive regulation of MAPK cascade, and inflammatory responses to be activated by HMZ treatment (Fig. 2b). KEGG analysis revealed distinct regulated pathways upon HMZ exposure, which include malaria, (p=1.65 x 10-7, FDR=2.59 x 10-6), MAPK signalling pathway (p=1.21 x 10-9, FDR=2.84 x 10-8), pathways of neurodegeneration (p=0.0325, FDR=0.0588) and Alzheimer disease (p=0.0446, FDR=0.0716) as upregulated pathways, and p53 signalling pathway (p=0.0069, FDR=0.0143) and apoptosis (p=0.0358, FDR=0.0576) as downregulated pathways (Fig. 2c, (Supplementary File 1).

Following the observations from the secretome profile, we performed qRT-PCR analysis to confirm mRNA expression of selected chemo-and cytokines in neuronal cultures of both cell lines (UM51 and B4) from treated and control conditions. Increased *IL-8* and *TNF-α* mRNA expression were observed in both cell lines upon HMZ treatment, while *IL-6* was only up-regulated in UM51 neuronal cultures (Fig. 2b). Furthermore, immune-regulatory, and inflammation-associated chemokines such as monocyte chemoattractant protein-1 (MCP-1) and macrophage migration inhibitory factor (MIF) showed increased secretion upon HMZ treatment (Fig. 2a, Supplementary Fig. 2). Secretion of vascular endothelial growth factor (VEGF) and platelet derived growth factor subunit B (PDGFB), both belonging to the same protein family and reported to be associated with inflammation, were also enhanced ^7^ (Fig. 2a). Additionally, enhanced mRNA expression and protein secretion of brain derived neurotrophic factor (BDNF) was observed. BDNF has been reported to be involved in modulating neuro-inflammation by providing neuroprotection^16^ (Fig. 2a, 2e). Next, we analyzed mRNA expression of selected genes associated with the mentioned pathways. Both UM51 and B4 neuronal cultures showed similar trends upon HMZ treatment in the neuronal cultures with an increase in *BDNF, C-C Motif Chemokine Ligand 2* (*CCL2*) expression, while both indicated a decrease in *VEGFA* and *PDGFB* expression (Fig. 2e). However, the decreased mRNA expression and increased protein secretion of VEGF and PDGFB point at additional factors besides transcription playing a role in the secretion of these proteins^17^. *CCL2* and *PECAM1* encode for the MCP-1 and CD31 protein, respectively. Both CCL2 and PECAM1 showed upregulated mRNA expression in both UM51 and B4-derived neuronal cultures after HMZ treatment (Fig. 2e).

### HMZ exposure induces DNA damage in UM51-derived neuronal cultures

We performed immunocytochemistry (ICC) and western blot (WB) analysis to evaluate the expression of the DNA damage marker-γH2AX (Fig. 3a, 3b). Both cell lines showed a similar trend with increased γH2AX levels in the WB analyses after HMZ exposure compared to the corresponding controls (Fig.3b, 3c).

**Figure 3.**
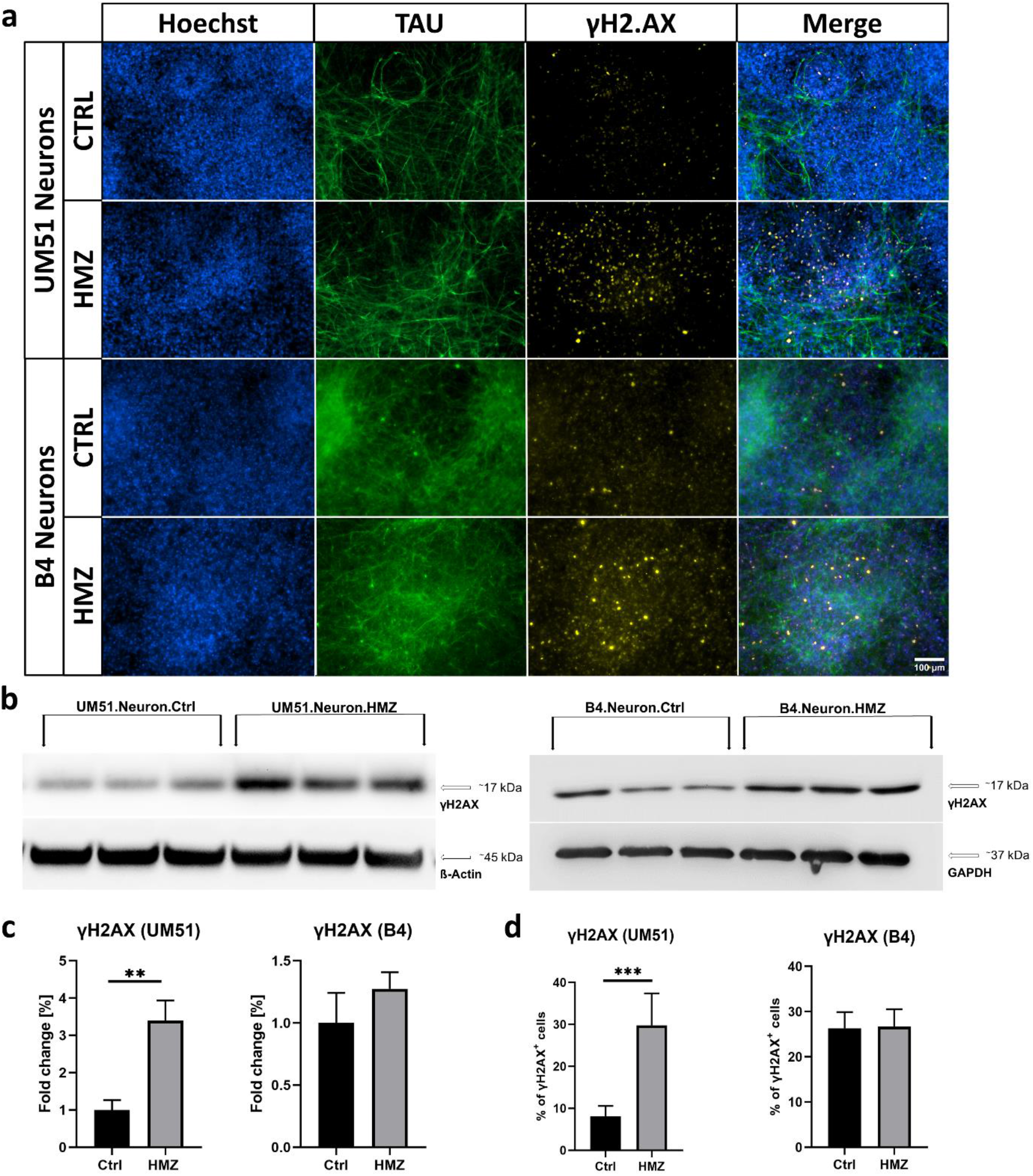
HMZ exposure induces DNA damage in iPSC-derived neuronal cultures. All depicted results were recorded in UM51- and B4-derived neuronal cultures after 48 h HMZ exposure and compared to their corresponding controls. **(a)** Representative ICC images showing TAU- and γH2AX-positive cells. Scale bar 100 μm. **(b)** WB analysis for γH2AX. **(c)** Bar graphs showing γH2AX protein expression in fold change. Values were normalized to ß-Actin and GAPDH (housekeeping gene) and subsequently to control samples. **(d)** Manual ICC quantification of γH2AX-positive nuclei. (c,d) n=3 for each condition, blots depict mean and error bars depict SD of all experiments. Asterisk (*) depicts significance, which is indicated by **p<0.01; ***p<0.001. Approximately six random fields from three separate differentiations for each condition were manually analysed (d).

Manual counting of nuclei in ICC staining showed 30% γH2AX-positive cells after 48 h of HMZ exposure in the UM51 neuronal culture, while the control condition consisted of 6% γH2AX positive cells (Fig. 3a, 3d). On the contrary, B4 neuronal cultures showed almost no change in ICC staining analysis (Fig. 3a, 3d). Aligning with the observation from ICC, in WB analysis UM51 neurons showed stronger response to HMZ treatment than the B4 neurons with a 3.4-fold increase in γH2AX expression, while the B4 neurons showed only a 1.3-fold increase compared to their corresponding controls (Fig. 3b, 3c).

### HMZ potentially activates p38 MAPK and not p53 signalling pathway in iPSC-derived neurons

As HMZ treatment on iPSC-derived neuronal cultures indicated p53 signalling pathway to be downregulated (Fig. 2d), and γH2AX levels were increased (Fig. 3), we investigated selected p53 signalling pathway-associated markers involved in DNA damage response and repair mechanisms (Supplementary Fig 3)^18-20^. First, we analysed mRNA expression of the DNA damage and repair related genes in both UM51 and B4-derived neuronal cultures. We observed p53 and MDM2 to be downregulated and ATM, ATR, CHEK1, CHEK2 to be significantly upregulated compared to the corresponding controls (Fig. 4a, 4b).

**Figure 4.**
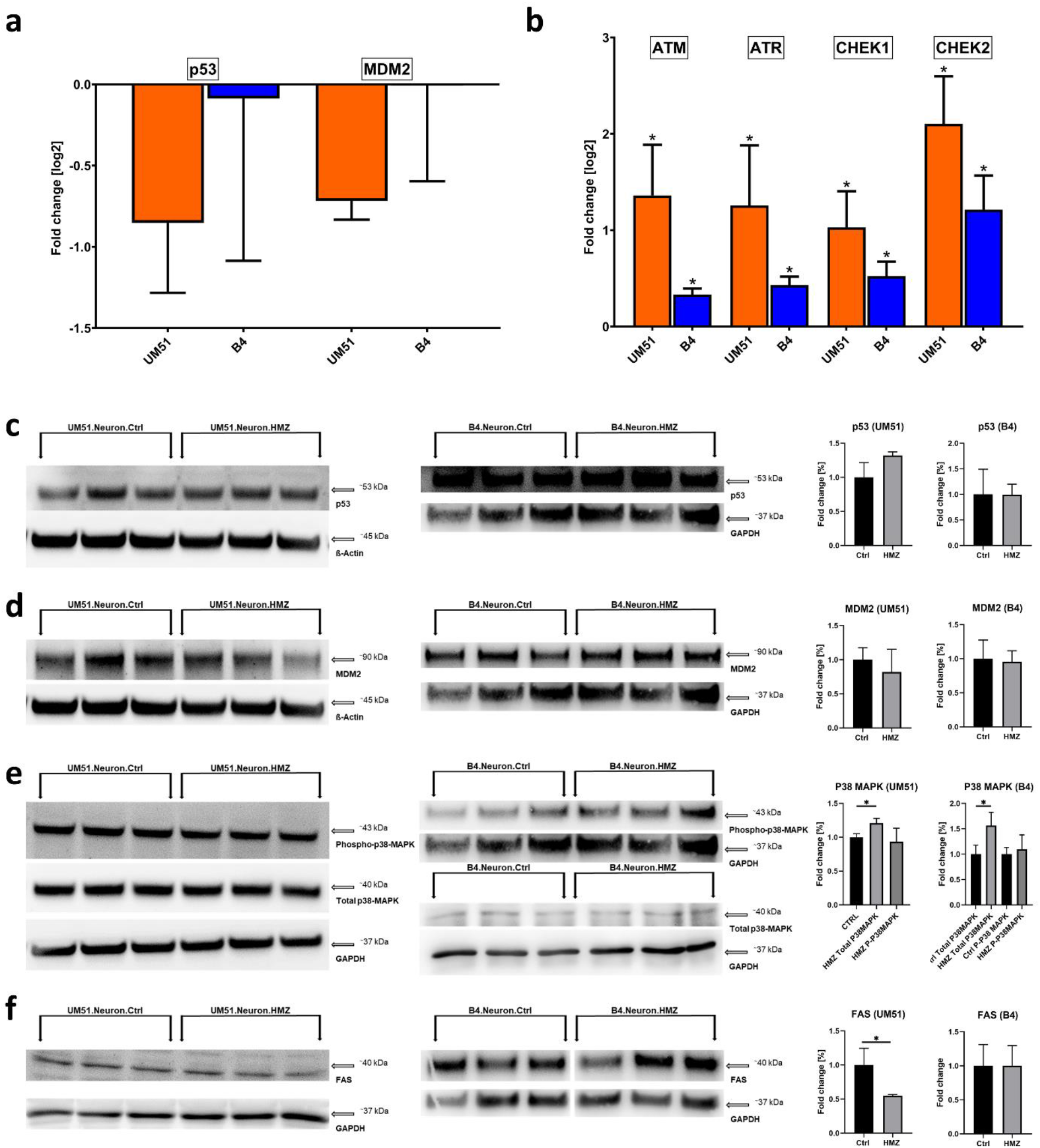
HMZ exposure activates DNA damage responses. All depicted results were performed in UM51- and B4-derived neuronal cultures after 48 h HMZ exposure and compared to their corresponding controls. **(a)** Relative mRNA expression analysis of *p53* and *MDM2*. **(b)** Relative mRNA expression analysis of *ATM, ATR, CHEK1* and *CHEK2*. **(c-f)** WB analyses and quantification of WB analyses for p53, MDM2, total p38, phospho-p38 and FAS. Values were normalized to ß-Actin and GAPDH and subsequently to control samples. (a,b) UM51 n=3; B4 Ctrl n=2; B4 HMZ n=3 (c-f) n=3 for each condition. (a-f) Blots depict mean and error bars depict SD of all experiments. Asterisk (*) depicts significance, which is indicated by *p<0.05.

In the WB analysis, p53 protein expression of the HMZ treated UM51 sample showed a slight increase, while the other P53 signalling-associated markers MDM2 (in both cell lines) and FAS (in UM51) showed a downregulated expression (Fig. 4c, 4d, 4f). All observations regarding these mentioned p53 signalling pathway-associated gene and protein expressions indicate a more vigorous response of the UM51 neuronal cultures upon HMZ exposure compared to B4 neuronal cultures (Fig. 4). This observation aligns with the observed higher DNA damage in the UM51 neuronal cultures compared to B4 (Fig. 3). Further experiments with immortalized HMC3 microglia-like cells also followed a similar trend with regards to p53 signalling pathway-associated markers and γH2AX expression after 48 h of HMZ exposure (Supplementary Fig. 4). Additionally, the change in total P38MAPK and phospho-P38MAPK protein expression might indicate MAPK signalling pathway activation after HMZ treatment in the iPSC-derived neuronal cultures (Fig. 4e, Fig. 2d).

## Discussion

An acute neurological complication of *P. falciparum* infection is CM, which is marked by neurological sequela in some survivors. The molecular mechanisms that underlie CM remain an enigma, which can partly be attributed to the lack of a suitable model that captures the clinical outcomes observed in patients. In this study, we employed neuronal cultures derived from iPSC lines obtained from two healthy ethnically diverse males and HMZ, the malaria toxin, as an *in-vitro* model of CM. TAU, MAP2 and β3-Tubulin-positive neuronal cultures were then stimulated with 20 μM HMZ for 48 h and deposition of the malarial toxin as brown crystals around neurons was observed. The deposition of HMZ as insoluble crystals into tissues is a common feature reported in both experimental and clinical malaria^21,22^. Indeed, the severity of falciparum infection is often correlated with the deposition of HMZ in organs^23^ and HMZ granules have been previously linked to thrombosed micro-vessels in fatal CM induced hemorrhage^24^.

Central to the progression of CM is neuro-inflammation. Notable inflammatory markers reported in CM include IL-1β, IL-6, IL-8, IFN-γ, TNF-α, PECAM-1, VCAM-1 and ICAM-1^25-27^. In this investigation, a secretome analysis using neuronal cell culture supernatant from UM51 cell line after HMZ treatment was conducted. Pearson’s heatmap analysis revealed an elevated IL-1β, IL-8 (partly), IL-16 and IFN-γ secretion. On the other hand, the levels of TNF-α, IL-6, IL-4 and IL-13 were lower in the HMZ treated cells. Interestingly, there was also a surge in the inflammatory-associated proteins: PDGFB, MIF, MCP-1 and VCAM-1. The increase in expression of these molecules is indicative of an inflammatory environment driven predominantly by a T helper type 1 (Th1) phenotype. Previous studies in clinical malaria have demonstrated that patients with CM had higher levels of pro-inflammatory Th1 cytokines as compared to severe malarial anemia patients^28,29^, which could account in part for the acute injuries observed in CM. In addition, these Th1 cytokines and inflammation associated proteins may be crucial to the development of neurocognitive and psychiatric disorders in CM patients. For instance, IL-8, IL-16 and MCP-1 have been implicated in neurocognitive symptoms in individuals with depression and anxiety^30^, and IL-1β and IL-16 were associated with psychiatric symptoms in schizophrenia patients^31^. Again, in neurodegenerative research, IL-16 was reportedly increased in patients with cerebrovascular dementias and AD^32^. Although IL-16 has been associated with psychiatric disorders and AD, this is the first study to implicate the cytokine in an experimental CM model. The cytokine was found to be significantly higher in asymptomatic falciparum patients when compared to individuals infected with Chikungunya virus^33^. Hitherto, there have been no reports on this cytokine in CM studies involving humans, which necessitates further research to understand the relationship between IL-16 and neurological impairment in CM patients. The reduced concentrations of IL-6 and TNF-α could be attributed to a delayed protein synthesis at the time of the assay. The pathogenesis of CM is characterized by persistent up-regulation of Th1 cytokines and inadequate production of anti-inflammatory cytokines including IL-4 and IL-13, which could possibly account for the observed lower levels of IL-4 and IL-13^34^. Instead of anti-inflammatory cytokines, the neurotrophic factor BDNF was elevated, which indicates a cellular response to remedy the highly inflammatory environment. In a study involving Ugandan children, McDonald et al., reported that lower BDNF levels were associated with poor prognosis in patients with CM, whereas higher levels of BDNF resulted in faster recovery from coma^35^. Using iPSC-derived brain organoids, Harbuzariu et al., discovered elevated levels of BDNF in response to heme injury^11^. Moreover, reduced BDNF expression has been linked to inflammation-induced apoptosis in neurons^36^, signifying a protective role of this neurotrophin against neuronal damage.

Metascape-based analysis data, obtained from up- and down-regulated proteins identified in the secretome screening, revealed an enrichment of proteins associated with malaria, MAPK cascade and inflammatory processes. The results confirm that HMZ is capable of activating cellular processes involved in malaria infection, even in the absence of the parasite. It also implies that the molecules elevated in our *in-vitro* CM model are relevant in malaria pathogenesis. With reference to the KEGG analysis, there was an up-regulation of pathways associated with malaria, MAPK signalling, neurodegeneration, and AD, whereas p53 signalling and apoptosis pathways were down-regulated. The up-regulated pathways support data from previous studies that identified genes related to neurodegenerative and AD pathways in CM patients^4,37,38^. This signifies that CM and neurodegenerative diseases, especially AD, share similar cellular mechanisms with a common denominator being inflammation. In contrast, the down-regulated p53 and apoptotic pathways can be attributed to the significant concentration of BDNF in our assay. Indeed, it has been confirmed that BDNF inhibits pro-apoptotic signalling following brain injury^11^. Saba et al., confirmed that BDNF significantly decreased apoptosis in serum deprived astrocytes by reducing p53 and active Caspase-3 expression^39^, hence the down-regulation of these pathways in our experiment.

Subsequent to secretome screening, the mRNA expression levels of pro-inflammatory cytokines associated with severe CM (TNF-α, IL-6, IL-8)^28^ but had a reduced/marginal protein expression, were examined in both UM51 and B4 neuronal cultures. Our analysis revealed significant upregulation of *TNFA* and *IL-8* in both neuronal cultures, and *IL-6* in only the UM51 neurons. The outcome demonstrates the phenomenon of poor correlation between mRNA and protein synthesis^40^, which explains the lower concentration of these cytokines in the secretome analysis, though differences in mRNA levels were significant.

Additionally, we examined the mRNA expression levels of genes associated with the up-regulated KEGG pathways in both cell lines namely: *BDNF, CCL2* (MCP-1), *PECAM1, PDGFB* and *VEGF*. With the exception of the angiogenic factors, *PDGFB* and *VEGF*, we saw an increase in mRNA expression of the other molecules. It is noteworthy that PDGFB and VEGF were elevated in the secretome analysis, therefore the lower mRNA levels once again demonstrate a poor correlation between mRNA and protein synthesis. Collectively, these experiments provide evidence that CCL2, PECAM1, PDGF and VEGF, which are reported to modify cell junctions, increase endothelial permeability and enhance leukocyte trans-endothelial migration in inflammation^41-44^ are key elements to HMZ induced cellular injury in CM. Ultimately, the significant expression of these factors represents inflammation and immune cell extravasation during CM pathogenesis.

A consequence of continuous inflammation is DNA damage. Therefore, in our successive experiments, we measured γH2AX, a reliable marker for DNA double strand breaks^45^ in both UM51 and B4 neuronal cultures. Additionally, we examined a selection of molecules involved in DNA damage and repair checkpoints including ATM, ATR, CHEK1 and CHEK2. Our analyses revealed a significant increase in γH2AX level, representing DNA damage in the UM51 neuronal cultures and only a slight increase in B4. This result is the first to confirm that HMZ induces DNA damage in neurons. The DNA checkpoint investigation revealed significant expression of *ATM, ATR, CHEK1* and *CHEK2* in both cultures, but more evident in UM51. Next, we measured downstream targets of the damage response pathway, p53 and p38 MAPK. We found the expression of *p53* transcripts to be lower in both cultures, but the expression of p53 protein to be slightly increased in UM51. We also detected lower protein levels of MDM2 (a negative regulator of p53) and FAS, a downstream target of p53. However, except for FAS, the changes were minimal. These results indicate that p53 pathway was not involved in the stress response in our experiments. On the contrary, we observed an up-regulation of total p38 MAPK in both cell lines, though there was no significantly higher expression of phosphorylated p38 MAPK. This is an indication that p38 pathway was activated in response to stress in our study. Previous research has described an activation of the p38 MAPK pathway in response to DNA damage through ATM, when p53 pathway was deficient or compromised^46,47^. Moreover, p38 MAPK was found to mediate the release of lysozyme and upregulate other pro-inflammatory molecules in an experiment involving human adherent monocytes and natural HMZ^48^. Again, p38 MAPK was up-regulated in peripheral blood mononuclear cells of patients infected with *P. falciparum*^49^. These observations suggest a vital role of p38 MAPK in facilitating cellular response to *P. falciparum* and HMZ.

As a subsequent goal, we examined the effect of HMZ on HMC3 microglia cells. There was an increase in the expression of known microglial activation markers-triggering receptor expressed on myeloid cells 2 (*TREM2*), ionized calcium binding adaptor molecule 1 (*IBA1*) and *CD45*. Compared to the control group, the level of γH2AX in HMZ treated microglia cells was higher though not significant. With the exception of ATR and CHEK1, there was an up-regulation of the other DNA damage response molecules (ATM, CHEK2) in HMC3 cells. Once more, there was no significant increase in p53 levels in HMZ treated microglia cells. Thus, neurons and microglia cells show a similar response pattern to HMZ stimulation.

A limitation of this study is the lack of complete cellular components characteristic of the brain, which comprise of neurons, glial cells, microglia, oligodendrocytes and vascular units. This would have provided the complexity of varying responses of the different cell types to HMZ exposure in CM pathology. Future experiments using brain organoids could be a suitable approach in this regard.

## Conclusion

This study aimed at identifying molecules and pathways that mediate neuronal response to HMZ. Additional to the well-established cytokines identified in CM, we discovered IL-16, which has been implicated in other neurological diseases but not in CM. We also provide evidence for the first time that HMZ induces DNA damage in neurons. In summary, this study has demonstrated that HMZ can potently induce inflammatory responses in a comparable manner to clinical falciparum malaria. In addition, iPSC-derived neuronal cultures may provide a robust model for CM research. Also, given the increasing evidence of a link between neurodegenerative diseases and CM, further studies are required to validate this assertion, specifically with regards to the molecules that drive the pathogenesis and the phenotype of these diseases.

## Supporting information

Supplementary figures, tables, files and whole western blot membranes

## Methods and Materials

### Cell culture

#### iPSCs culture and NPC generation

Two healthy male-derived iPSC lines were used in this study. The UM51-hiPSC line was derived from SIX2-positive renal progenitor cells isolated from the urine of a 51-year-old healthy male of African origin (UM51)^14^. The other iPSC line (B4) was generated from healthy human fetal foreskin fibroblasts (HFF1, ATCC, #ATCCSCRC-1041, http://www.atcc.org)^15^ (Supplementary Table S1). iPSCs were plated on Matrigel (Corning, New York, NY, USA)-coated culture dishes using mTeSR Plus medium (StemCell Technologies, Vancouver, Canada). Cultures were routinely tested for mycoplasma contamination. Cells were passaged by dissociation into small aggregates with ReLeSR (StemCell Technologies, Vancouver, Canada) every 5–7 days and split in a 1:5 ratio into fresh Matrigel-coated dishes. For neural progenitor cell (NPC) generation, iPSCs were split as single cells using accutase (Life Technologies, Waltham, MA, USA). In brief, at day 0, iPSC-colonies were dissociated with warm (37°C) accutase, centrifuged at 200 rcf for 5 min, then resuspended in mTESR+ with 10μM of ROCK inhibitor. 20,000 cells in 100 μL cell suspension were seeded per well of a 96 u-bottom well plate (Low attachment, U bottom) to form embryoid bodies (EBs). Seeded iPSCs were spinned down at 110 rcf for 3 min and incubated at 37°C with 5% CO_2_ for 24h. At day 1, half of the medium volume was aspirated and replaced with neural induction medium (NIM) with 10μM of ROCK inhibitor. EBs were culture for 7 days (day 2-8) with NIM supplemented with 10 μM SB431542 and 5 μM Dorsomorphin (10μM of ROCK inhibitor until day 3). Medium was refreshed daily. On day 8, 20-30 EBs were plated per well of a Poly-Ornithine/Laminin (Sigma-Aldrich; Merck KGaA, Darmstadt, Germany) -coated 6-well plate to generate neural rosettes. EBs were cultured in neural differentiation medium (NDM) (Neurobasal A, 1% B27, 1% GlutaMAX, 1% P/S) supplemented with 20 ng/mL EGF and 20 ng/mL FGF2 and incubated at 37°C with 5% CO_2_ in static position. The medium was changed daily. At day 16, neural rosettes were selected using STEMdiff™ Neural Rosette Selection Reagent (StemCell Technologies, Vancouver, Canada) for 30 min-1 h at 37°. Warm accutase was used for 30 min at 37°C to dissociate the neural rosettes. NPCs were further cultured in Growth-Factor Reduced (GFR) Matrigel (Corning, New York, NY, USA)-coated plates with NDM supplemented with 20 ng/mL EGF and 20 ng/mL FGF2. Accutase was used for splitting the NPCs. In parallel, early passages of NPCs were frozen in Cryostore CS10 (StemCell Technologies) for later usage.

#### Neuronal differentiation and Hemozoin treatment

UM51- and B4 iPSC-derived NPCs were used for the neuronal differentiation. To initiate differentiation, NPCs were dissociated with warm accutase and plated on GFR Matrigel (Corning, New York, NY, USA)-coated plates. 500,000 cells/well in a 6-well plate and 80,000 cells/well in a 24 well plate were seeded to produce neuronal cultures. For cell adhesion, NDM supplemented with 20 ng/mL EGF and 20 ng/mL FGF2 was used for 24h, which was then changed to NDM supplemented with 20 ng/mL BDNF and 20 ng/mL NT3 for the rest of the differentiation (Day 2-16). Cells were cultured until day 16 and medium was changed every 2-3 days. At day 16, neuronal cultures were exposed to 20 μM HMZ for 48 h.

Hemozoin (InvivoGen #tlrl-hz) was dissolved in PBS (pH 7.4) to obtain a stock concentration of 4.06 mM (5mg/mL). To obtain a homogenous dispersion of hemozoin, the hemozoin-PBS suspension was sonicated for 5 minutes. For treatment, 20 μM hemozoin (HMZ) was added to the culture medium and incubated for 48 h.

### Cell viability assay

To ascertain the appropriate concentration, we performed resazurin assay/alamar blue assay on B4 iPSC-derived neuronal cultures by exposing those to 0, 5 μM, 10 μM, 20 μM, 50 μM, 100 μM, 200 μM HMZ for 48 h. In brief, day 16 neurons were treated with the mentioned HMZ concentrations continuously for 48 h. 2 h before reaching the analysis time point, resazurin solution was added and incubated at 37°C with 5% CO_2_. After 2 h, 100 μl supernatant was transferred to a 96-well flat bottom plate in triplicate for each condition. Fluorescence was recorded using a plate reader with a 560 nm excitation/ 590 nm emission filter set. For the resazurin solution, a stock of 0.15 mg/mL resazurin in PBS was prepared, which was further diluted 1:10 with PBS before use. 100 μl diluted resazurin solution was added per well. 3 wells containing culture medium and diluted resazurin were prepared for background subtraction and instrument gain adjustment.

Based on Resazurin measurements of neuronal culture treated with varying doses of HMZ, IC50 plots were generated within the R environment employing the packages dr4pl and ggplot2^50,51^ (Supplementary Fig. 1). The dr4pl method was parametrized to use a logistic model, the fitted curve was plotted using values from the model. Additionally, the IC50 was calculated via the logistic model in dr4pl.

### Immunocytochemistry

Cells were fixed in 4% paraformaldehyde (PFA) (Polysciences, Warrington, FL, USA) for 10 min at room temperature (RT). After washing with 3 times with PBS, cells were directly used for staining. Fixed cells were permeabilized for 10 min using 0.1% Triton X-100 in PBS+Glycine (30 mM Glycine) at RT. After washing once with PBS, unspecific binding sites were blocked for 2 h at RT with 3% BSA in PBS+Glycine. Primary antibodies were diluted in 3% BSA in PBS+Glycine and incubated overnight at 4°C (Supplementary Table S2). After washing three times with PBS, secondary antibodies diluted in 3% BSA/PBS/Glycine were added for 2 h and incubated at RT. Nuclei were stained with Hoechst 33258. Stained cells were imaged using a Zeiss fluorescence microscope (LSM 700). Individual channel images were processed and merged with ImageJ software version 1.53c (U. S. National Institutes of Health, Bethesda, Maryland, USA). Cell counting was performed using the “Cell counter” function of ImageJ.

### Secretome Analysis

#### Human XL Cytokine Array

After 48 h of HMZ treatment, the conditioned medium of the control and HMZ-treated neuronal cultures were stored at -20°C. Relative expression levels of 105 soluble human proteins and cytokines were determined using the Human XL Cytokine Array Kit from R&D Systems. The cytokine array was performed following the manufacturer’s guidelines. In brief, the membranes were blocked for 1 h on a rocking platform using the provided blocking buffer. Samples were prepared by diluting the desired quantity to a final volume of 1.5 mL with array buffer (array buffer 6). The sample mixtures were pipetted onto the blocked membranes and incubated overnight at 4°C on a rocking platform. Membranes were washed three times with washing buffer for 10 min each at RT. Then, the membranes were incubated with detection antibody cocktail for 1h at RT and washed three times. Afterwards, Streptavidin-HRP was added onto the membranes, and incubated for 30 min at RT. ECL detection reagent (Cytiva, Freiburg, Germany) was used to visualize bound proteins as spots, which were detected using a Fusion FX instrument (PeqLab, Erlangen, Germany) (Supplementary Fig. 2).

#### Image Analysis of Cytokine Arrays

Images of the hybridized cytokine assays were scanned and analyzed via the FIJI/ImageJ software^52^. The grid on the cytokine array was detected semi-automatically employing preprocessing via Gaussian blur (size 4) and local maxima finding. The csv file containing the local maxima detected by FIJI/ImageJ was imported into the R programming environment and based on it corners were found and the grid between them interpolated making use of positions found as local maxima. Quantification of the spots at the grid locations was performed by the FIJI Microarray Profile plugin by Bob Dougherty and Wayne Rasband (https://www.optinav.info/MicroArray_Profile.htm, accessed on 21 December 2022), using integrated densities. Cytokine names were taken from proprietary lists of the manufacturer (Proteome Profiler Array from R & D Systems, Human XL Cytokine Array Kit, Catalog Number ARY022B) and assigned to the quantified spots by their grid positions.

#### Cytokine Data Analysis

The follow-up-processing of the quantified spots within the R/Bioconductor environment started with the Robust Spline Normalization from the R/Bioconductor package lumi^53,54^. The function heatmap.2 from the gplots package was applied to draw heatmaps using Pearson correlation as similarity measure^55^. Based on this cluster analysis via heatmap.2 genes were split into two clusters associated with up- and down-regulation by HMZ for subsequent analysis.

### Metascape Analysis

Comprehensive functional analysis of the clustered gene ontology (GO) biological processes and pathways (KEGG pathways, Reactome Gene Sets, Canonical pathways, and CORUM) of gene-sets based on data derived from the Human XL Cytokine Array was performed using Metascape^56^ (http://metascape.org, accessed on 13 October 2023). Lists of up- and downregulated cytokines were treated as gene-sets for analytical purposes.

### Pathway analysis

Cytokines were subjected to cluster analysis via the heatmap.2 function from the R package gplots using Pearson correlation as similarity measure and the default setting of hclust as hierarchical clustering method^55^. Cytokines from the two resulting clusters of HMZ-induced up- and down-regulation were analyzed for over-represented KEGG pathways in the follow-up analysis^57^. KEGG pathways were downloaded from the KEGG website in February 2023. The R-builtin hypergeometric test was employed to determine the p-value indicating the significance of the over-represented pathways. Malaria and p53 signaling (KEGG) pathway, shown in the supplementary fig. 3 were taken respectively from https://www.genome.jp/kegg-bin/show_pathway?hsa05144 and https://www.genome.jp/pathway/hsa04115.

### Reverse Transcriptase PCR (RT-PCR)

Control and HMZ-treated cells were lysed for 15 min in Trizol. RNA was isolated using the Direct-zol™ RNA Isolation Kit (Zymo Research, Freiburg, Germany) according to the user’s manual, including the DNA digestion step. 500 ng of RNA was reverse transcribed using the TaqMan Reverse Transcription Kit (Applied Biosystems, Waltham, MA, USA). The primer sequences are shown in (Supplementary Table S3). Real-time PCRs were performed in technical and independent experiment triplicates (n=3) with Power Sybr Green Master Mix (Life Technologies, Darmstadt, Germany) on a VIIA7 (Life Technologies, Darmstadt, Germany) machine. Mean values were normalized to the ribosomal protein lateral stalk subunit P0 (RPLP0) and fold-change was calculated using the indicated controls. Changes in gene expression were presented in log2-scale. All values are depicted as mean with a 95% confidence interval (CI). Each singular value (Ctrl and HMZ) was compared to the mean of all corresponding control samples. The resulting values were used for statistical analysis. Statistical analysis was performed using Student′s unpaired two-sample t-tests. For ease of reading, only values of the treated condition were presented.

### Western Blotting

Total protein from UM51- and B4-derived neuronal cultures was isolated using RIPA buffer (Sigma-Aldrich Chemicals, Taufkirchen, Germany), supplemented with protease and phosphatase inhibitors (Roche, Mannheim, Germany). Afterwards, Pierce BCA Protein Assay Kit (Thermo Fisher, Waltham, MA, USA) was used to determine protein concentration of the samples. 20 μg of the heat-denatured protein lysate of each sample was loaded on a 4–12% SDS-PAGE and then transferred via wet blotting onto a 0.45 μm nitro-cellulose membrane (GE Healthcare, Solingen, Germany). After 1 h of blocking with 5% milk in TBST, the membranes were stained with anti-γH2AX, anti-p53, anti-MDM2, anti-P38MAPK, anti-phospho-P38MAPK, anti-FAS, anti-Caspase 3, and anti-cleaved caspase 3 antibodies in the appropriate buffer (Supplementary Table S2). Incubation with primary antibodies was performed overnight at 4°C.

After washing the membranes three times with TBST, secondary antibody incubation was performed for 2 h at RT followed by washing with TBST (Supplementary Table S2). Anti-β-Actin and anti-GAPDH (glyceralde-hyde-3-phosphate dehydrogenase) were used as housekeeping proteins to normalize protein expression. ECL Western Blotting Detection Reagents (Cytiva, Freiburg, Germany) were used to visualize the stained protein bands and then detected in a Fusion FX instrument (PeqLab, Erlangen, Germany). Band intensity quantification and analysis was performed with Fusion Capt Advance software FX7 16.08 (PeqLab, Erlangen, Germany).

### Statistical Analysis

All obtained results were represented as mean ± standard deviation. Statistical analysis (Student′s unpaired two-samples t-test) was performed utilizing GraphPad Prism v.8.0.2 (GraphPad Software, Boston, USA). A p-value of <0.05 was considered as statistically significant during analysis.

## Acknowledgment

The authors are grateful to Kanehisa Laboratories for the permission to use their figures for Malaria and p53 signalling path for our figures. The figures were taken from https://www.genome.jp/kegg/

## Author contributions

A.I.P. and L.P.S. designed and performed the experiments, processed and analyzed the data, and wrote and edited the manuscript. W.W. performed the bioinformatic analysis and data curation, helped with the figures, wrote the bioinformatic section in the Methods and Materials, and edited the manuscript. A.A.K. conceptualized the work, wrote and edited the manuscript. J.A. conceptualized and designed the work, edited the manuscript, acquired funding, and supervised the study. All authors have read and agreed to the submitted version of the manuscript.

## Data availability statement

Datasets resulting from the cytokine assay analyses in this study are available in the supplementary materials (Supplementary File 1, 2) data of this work.

