## Supplementary figures, tables, files and whole western blot membranes for "Hemozoin induces Malaria via activation of DNA damage, p38 MAPK and Neurodegenerative Pathways in Human iPSC-derived Neuronal Model of Cerebral Malaria": Supplementary Figures.pdf

**Supplementary Fig. 1:**

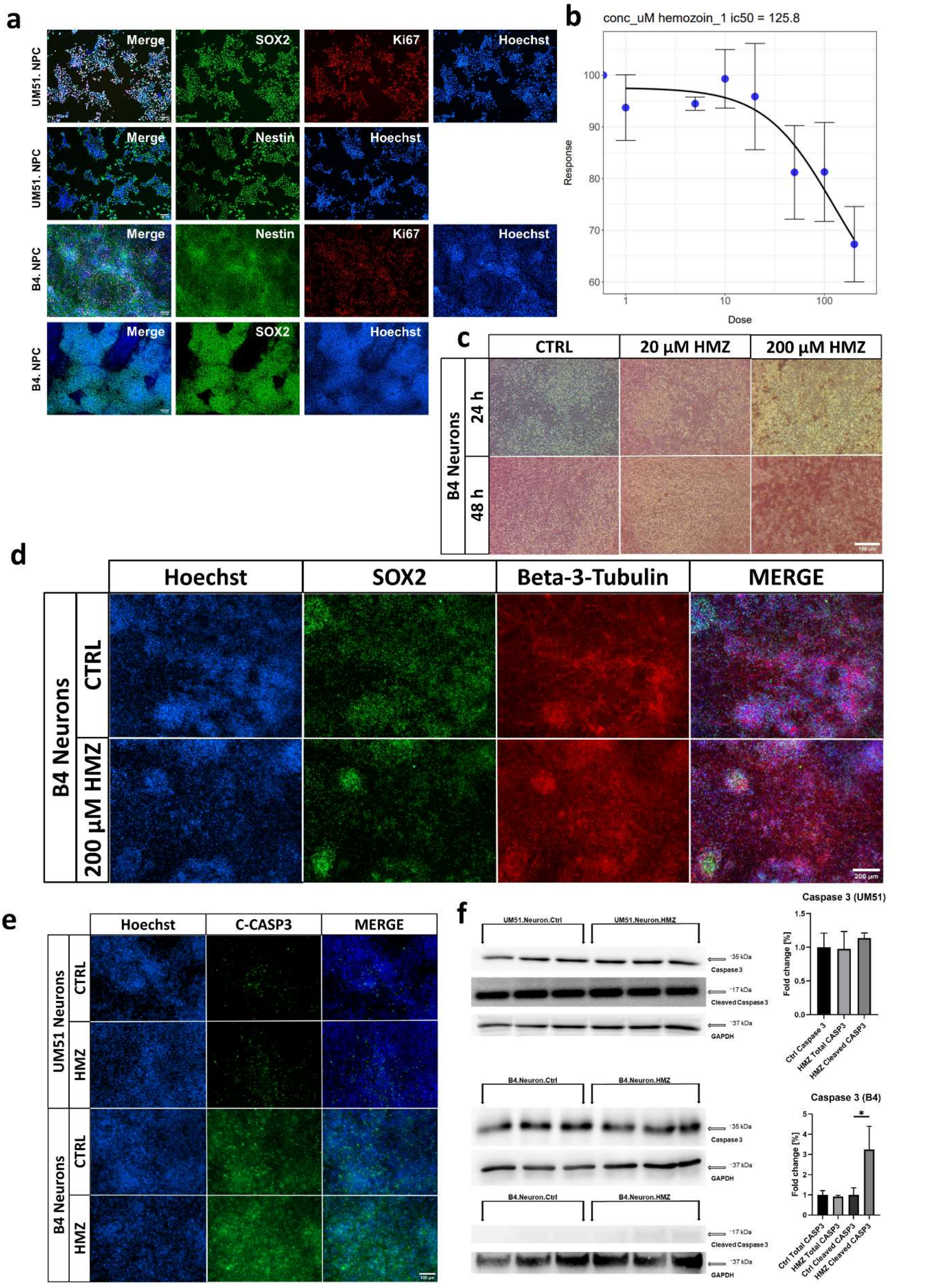


**Supplementary Figure 1. Preliminary and optimal-dose experiments for HMZ treatments (a)** Representative ICC images of Nestin-, SOX2- and KI67-positive cells in cultured neural progenitor cells. **(b)** Dose-response curve of B4-derived neuronal cultures treated with 0, 1, 5, 10, 20, 50, 100 and 200 µM HMZ. **(c)** Representative brightfield images of control, 20 µM and 200 µM HMZ-treated B4 iPSC-derived neuronal cultures at 24h and 48h of exposure. Scale bar 100 µm. **(d)** Representative ICC images of SOX2- and β3-Tubulin-positive cells in B4-derived neuronal networks after 48h of 200 µM HMZ exposure in comparison to control. Scale bar 200 µm. **(e)** Representative ICC images of Cleaved Caspase 3-positive cells in UM51- and B4-derived neuronal cultures after 48h exposure to 20 µM HMZ in comparison to control. Scale bar 100 µm. **(f)** WB analyses and quantification of WB analyses for total Caspase 3 and Cleaved Caspase 3 in in UM51- and B4-derived neuronal cultures after 48h HMZ exposure in comparison to control. n=3 for each condition, blots depict mean and error bars depict SD of all experiments. Asterisk (*) depicts significance, which is indicated by *p<0.05.

**Supplementary Fig. 2:**

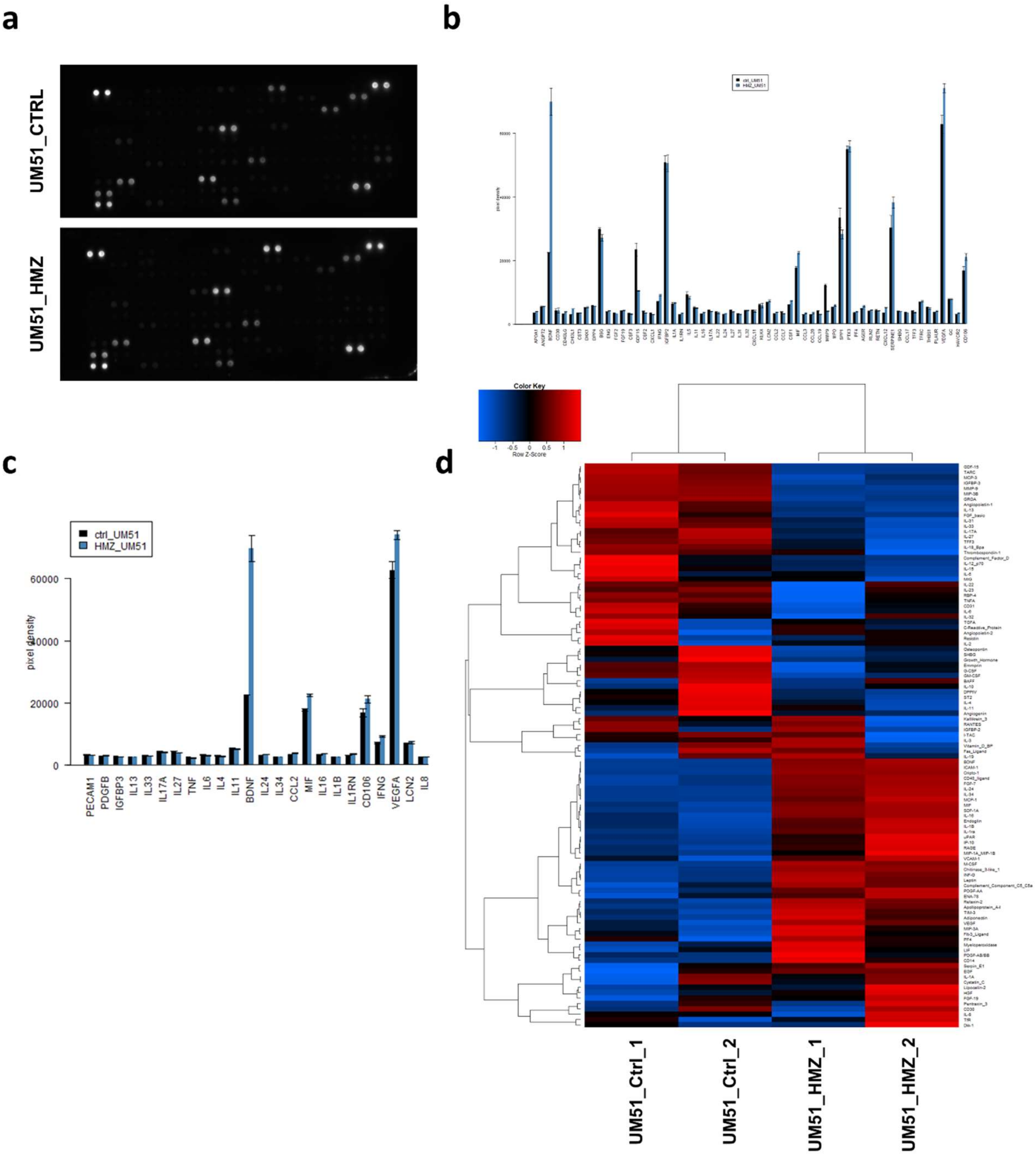


**Supplementary Figure 2. UM51 neuronal culture cytokine array. (a)** Cytokine array blots utilized for detecting altered secretomes in UM51 neuronal cultures after 48h 20 µM HMZ exposure in comparison to control. **(b)** Multibar plot showing all chemo- and cytokines with pixel density values over background pixel density. **(c)** Multibar plot for selected chemo- and cytokines showcased as heatmap in Figure 2a. **(d)** Heatmap using Pearson correlation as similarity measure depicting all examined chemo- and cytokines. Error bars denote the standard error of the mean (b,c).

### Supplementary Fig. 3:

**a**

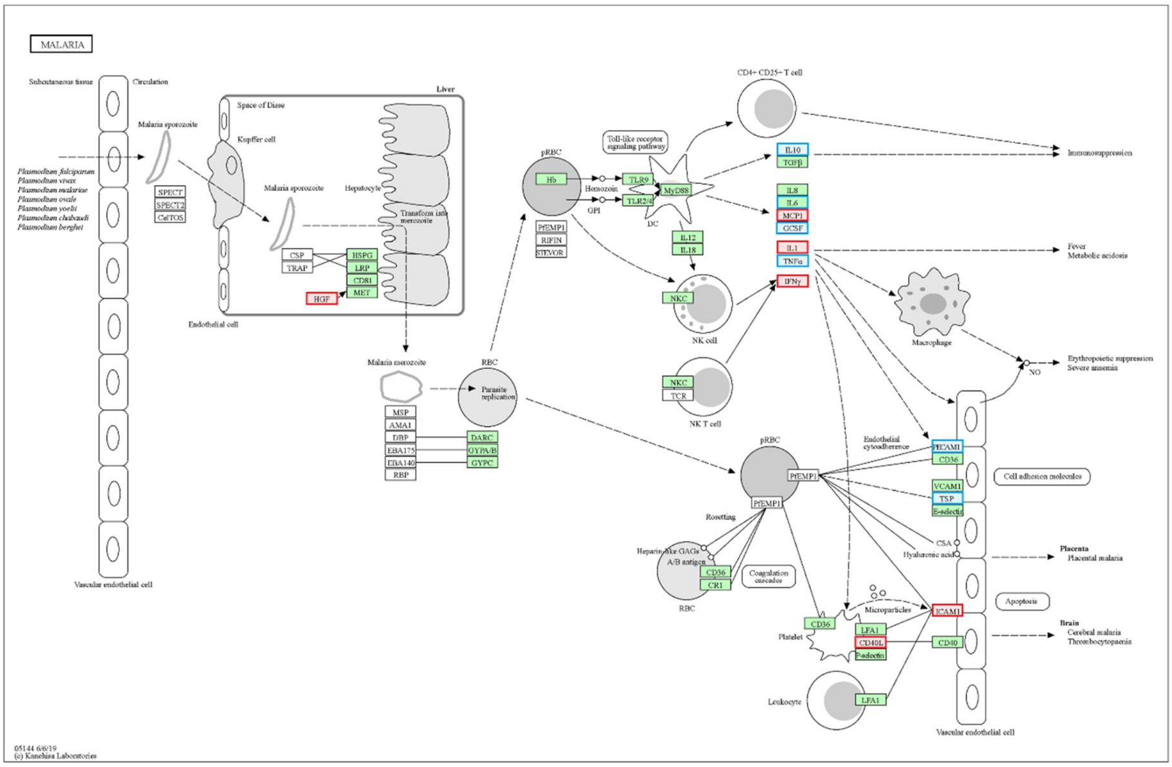


**b**

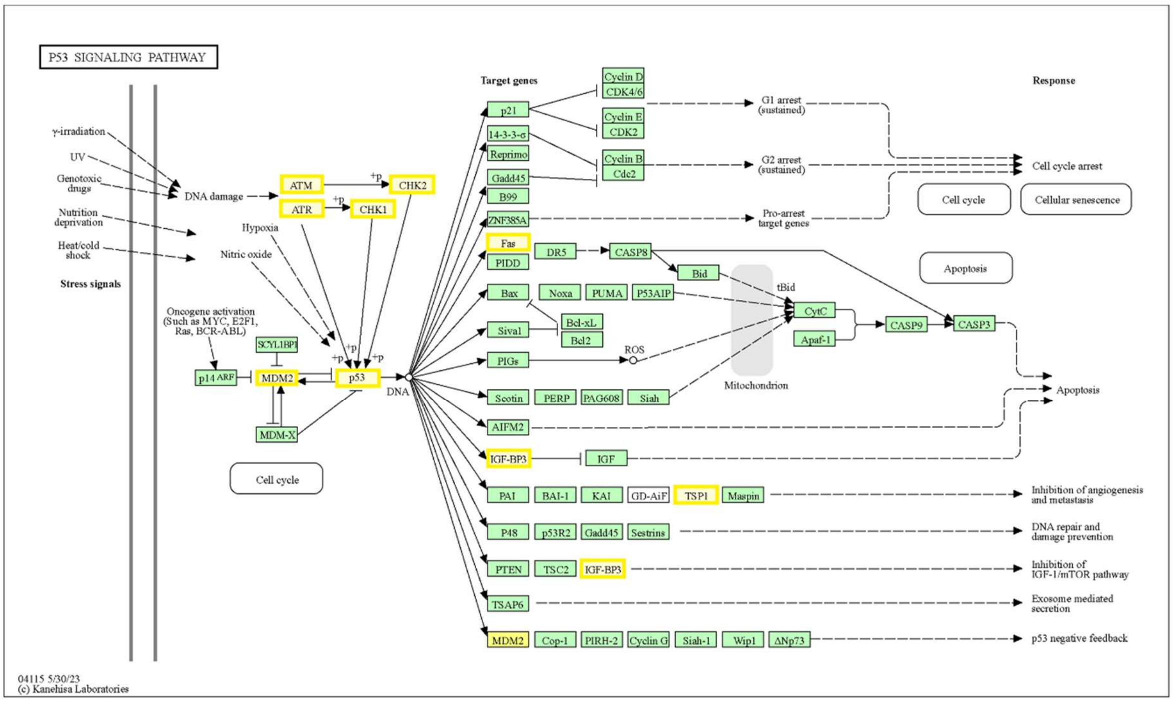


**Supplementary Figure 3. Analysis of the KEGG pathways - Malaria and p53 signalling pathways in iPSC-derived neuronal cultures. (a)** Schematic of the KEGG pathway Malaria. Cytokines upregulated in the secretome of UM51 neuronal cultures are marked in red, downregulated in blue. **(b)** Schematic of the KEGG pathway p53 signalling pathway. Regulated genes are highlighted in yellow, and non-regulated in grey.

**Supplementary Fig. 4:**

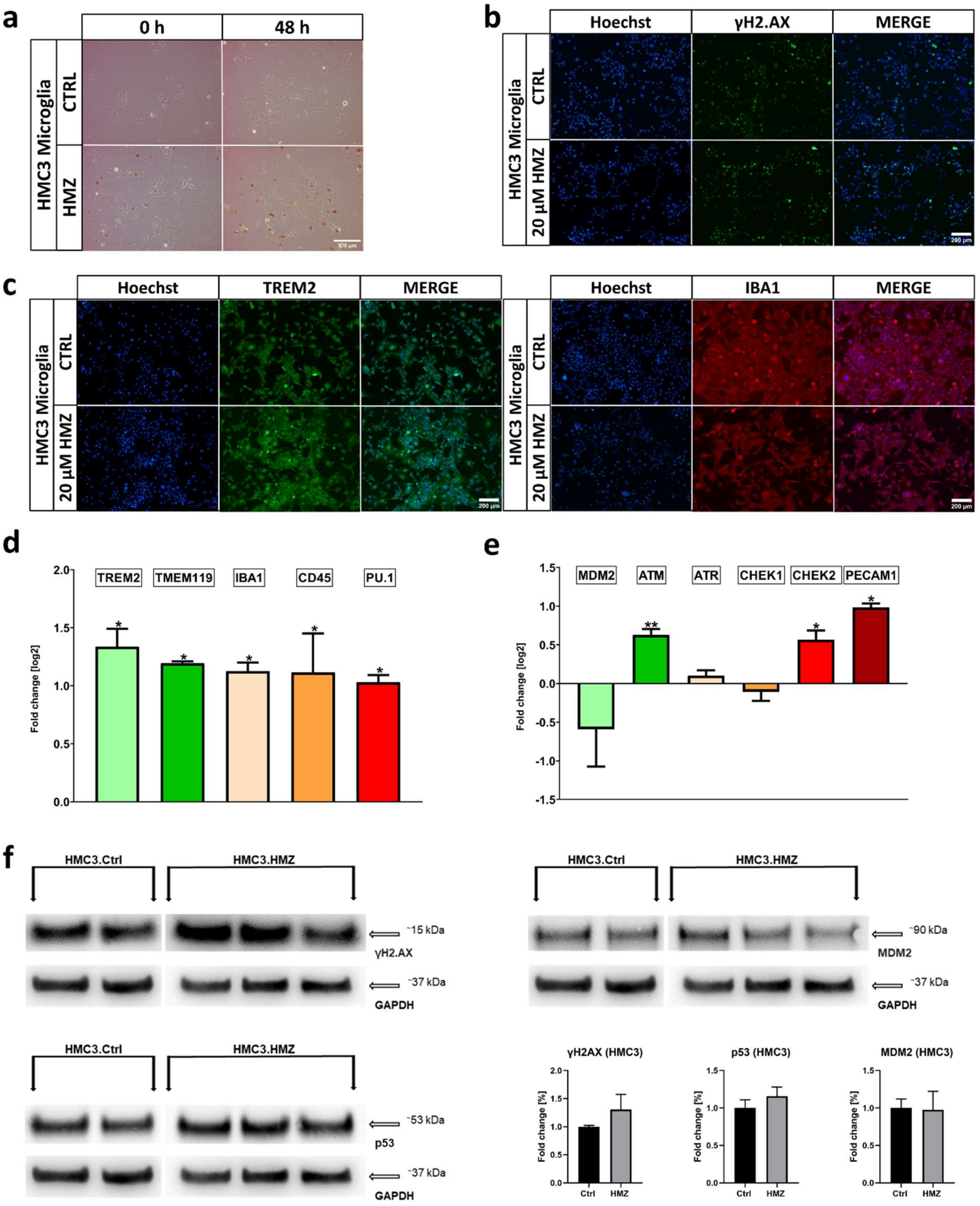


**Supplementary Figure 4. HMC3 cells show signs of microglial activation and activation of DNA damage response. (a)** Representative bright field images of control and 20 µM HMZ-treated HMC3 cells at 0h and 48h of exposure. Scale bar 100 µm. **(b,c)** Representative ICC images of γH2AX-, TREM2- and IBA1-positive HMC3 cells after 48h HMZ exposure in comparison to control. Scale bar 200 µm. **(d)** Relative mRNA expression analysis of *TREM2, TMEM119, IBA1, CD45* and *PU.1* in HMC3 cells after 48h HMZ exposure in comparison to control. **(e)** Relative mRNA expression analysis of *MDM2, ATM, ATR, CHEK1, CHEK2* and *PECAM1* in HMC3 cells after 48h HMZ exposure in comparison to control. **(f)** WB analyses and quantification of WB analyses for γH2AX, MDM2 and p53 after 48h HMZ exposure in comparison to control. Values were normalized to GAPDH and subsequently to control samples. Ctrl n=2; HMZ n=3. (d,e) n=3; blots depict mean and error bars depict SD of all experiments. Asterisk (*) depicts significance, which is indicated by *p<0.05; **p<0.01.
