## Supplementary figures, tables, files and whole western blot membranes for "Hemozoin induces Malaria via activation of DNA damage, p38 MAPK and Neurodegenerative Pathways in Human iPSC-derived Neuronal Model of Cerebral Malaria": Whole blots.pdf

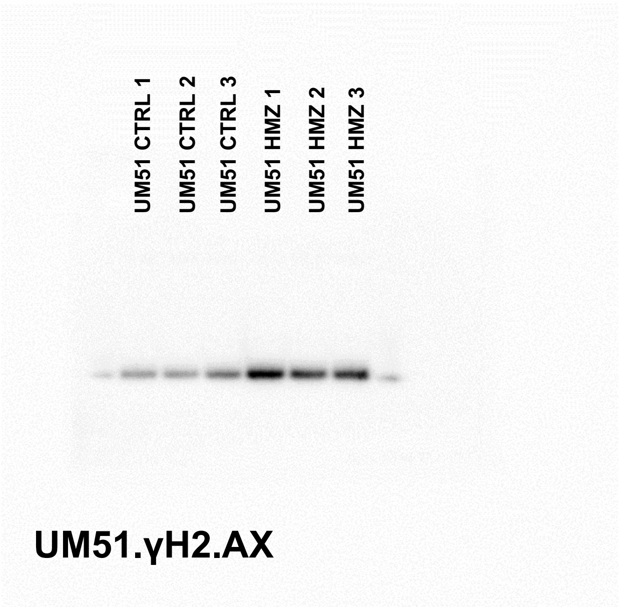


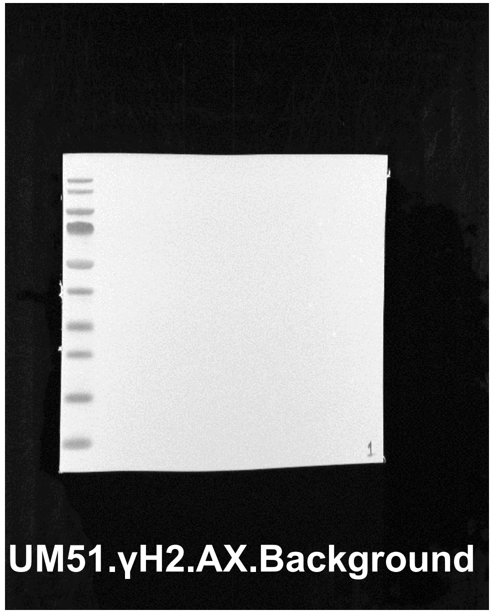


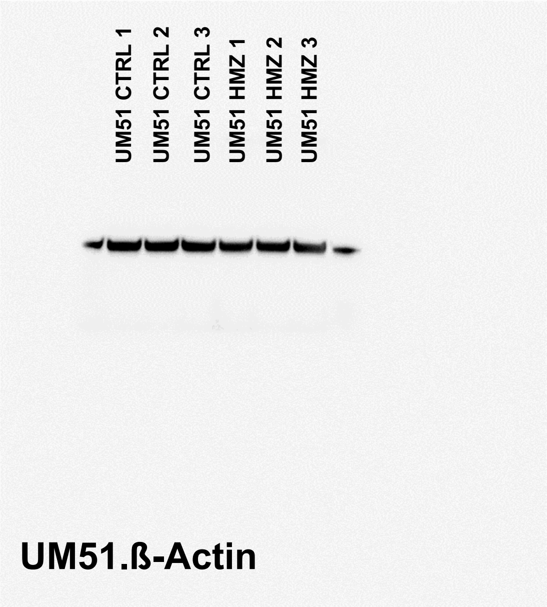


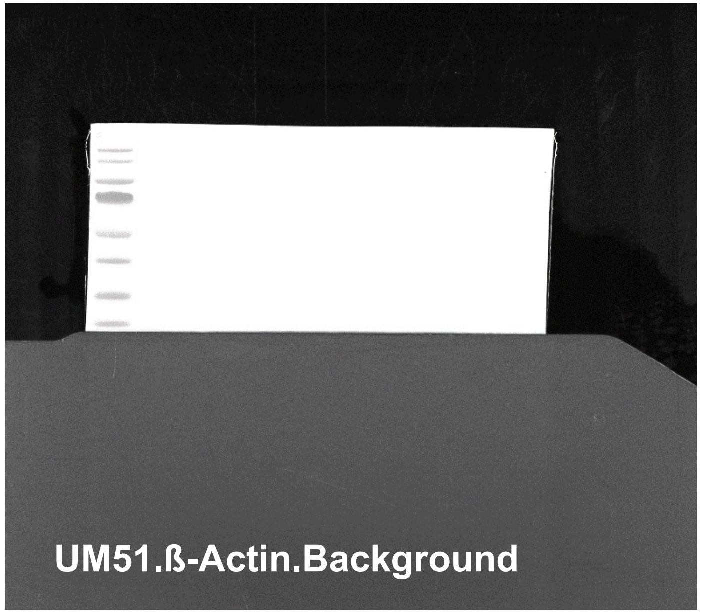


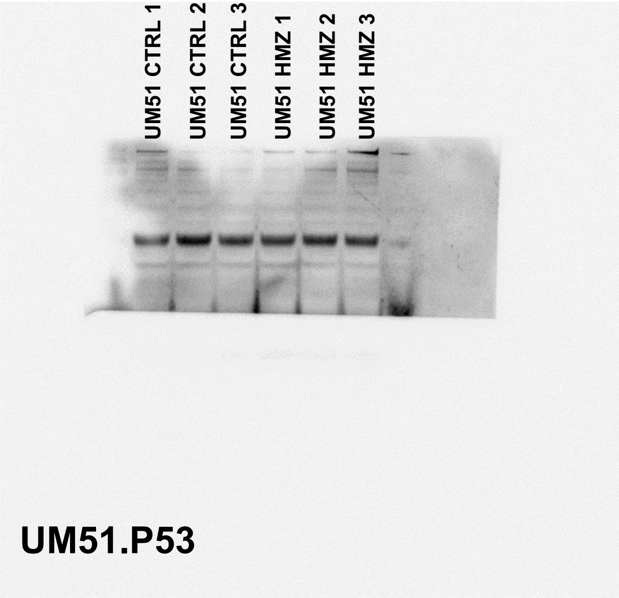


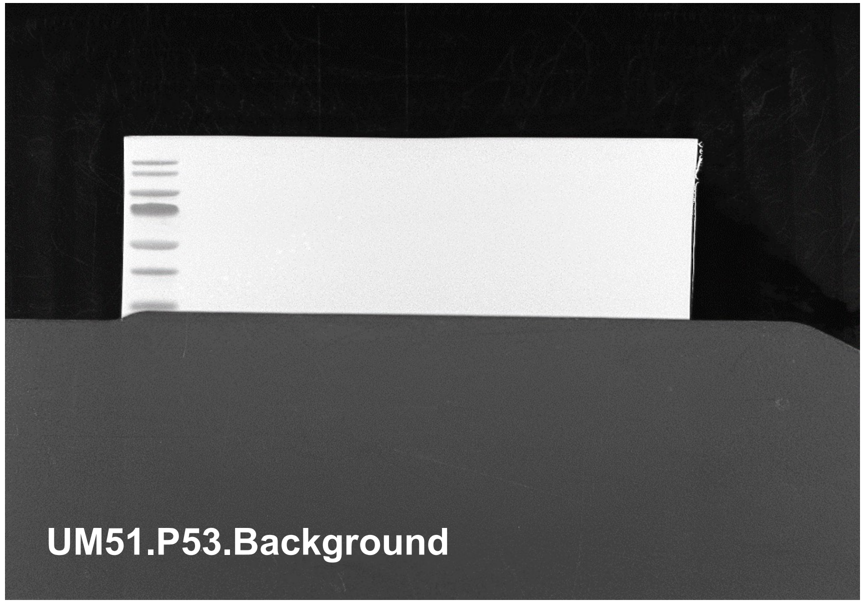


The whole blot membranes for main figure 3,4. The stainings were done on the same membrane for γH2AX and P53.

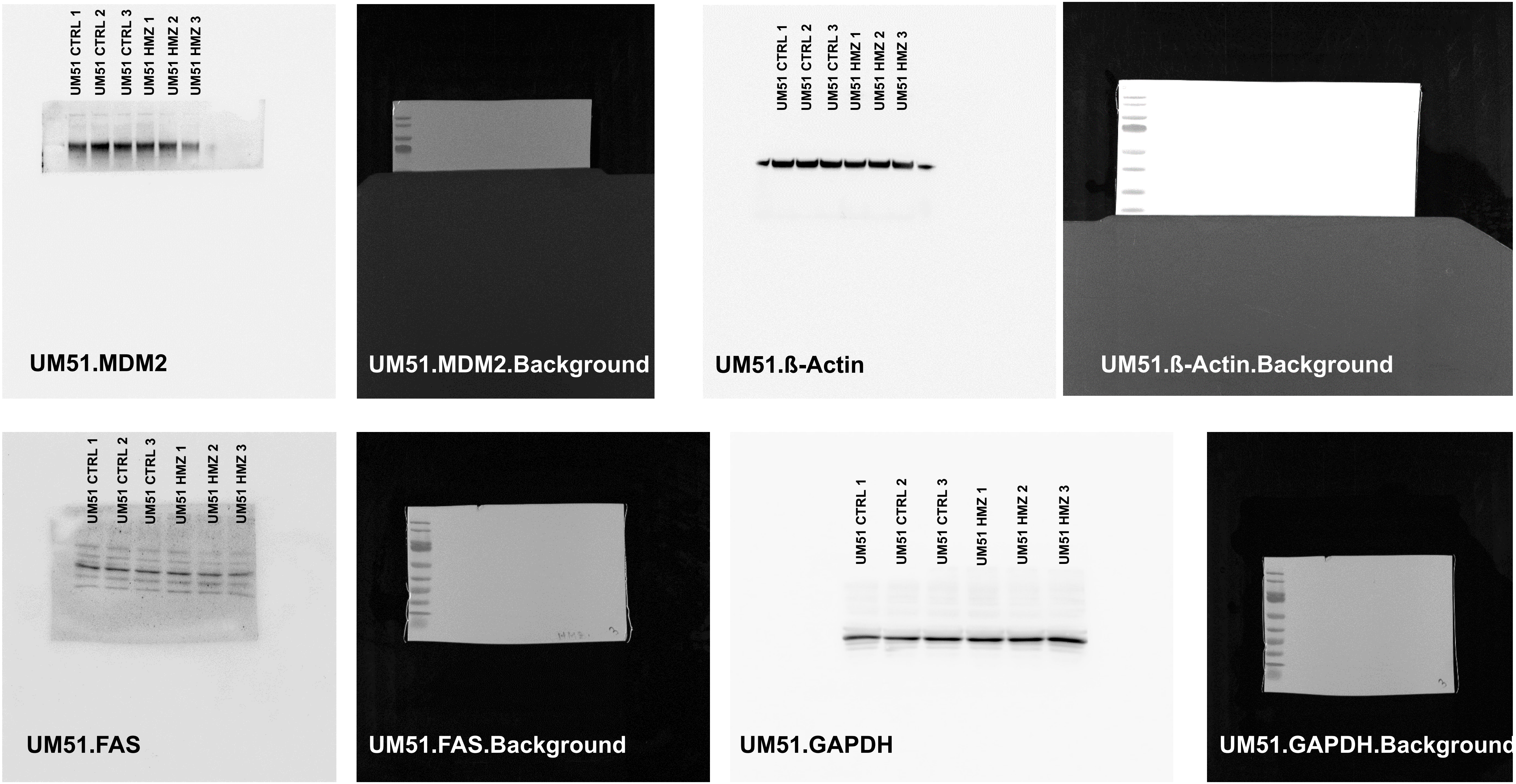


The whole blot membranes for main figure 4.

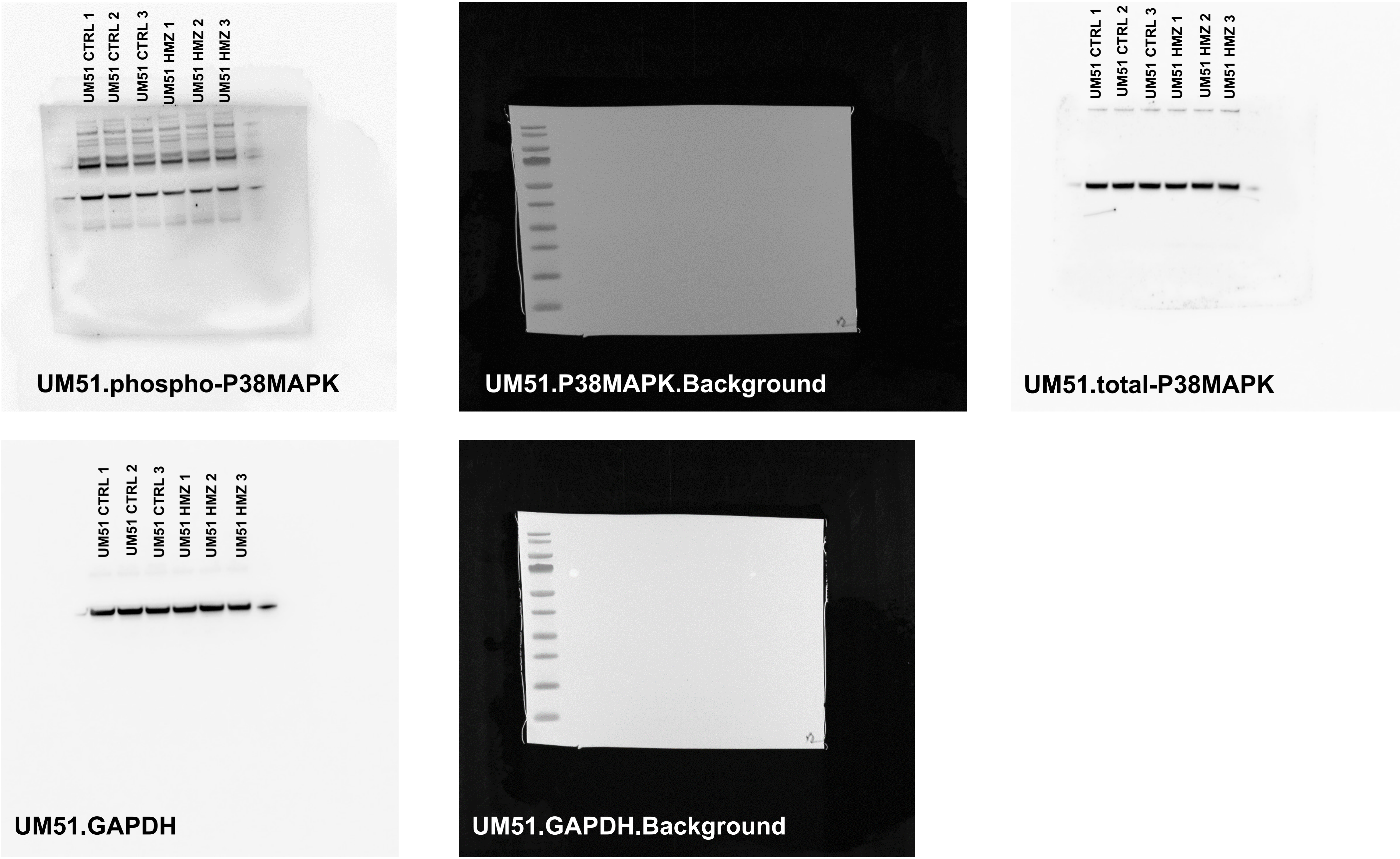


The whole blot membranes for main figure 4.

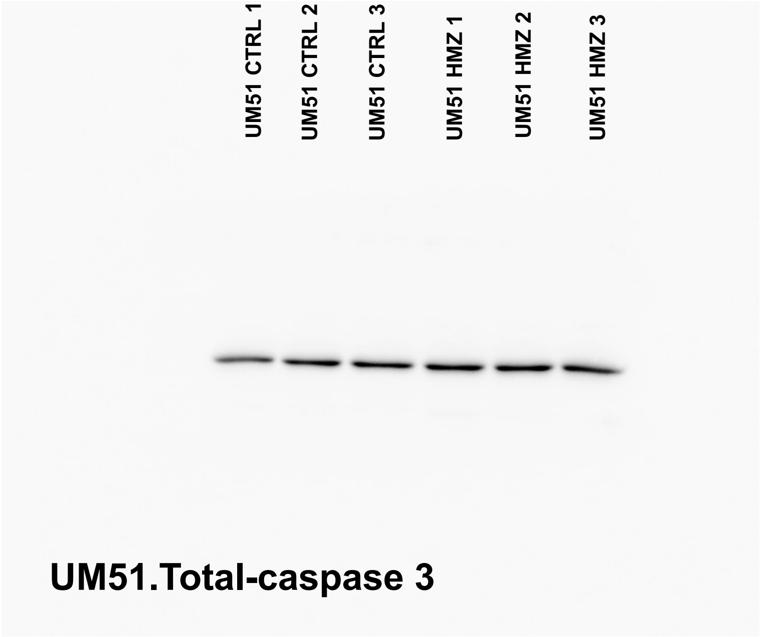


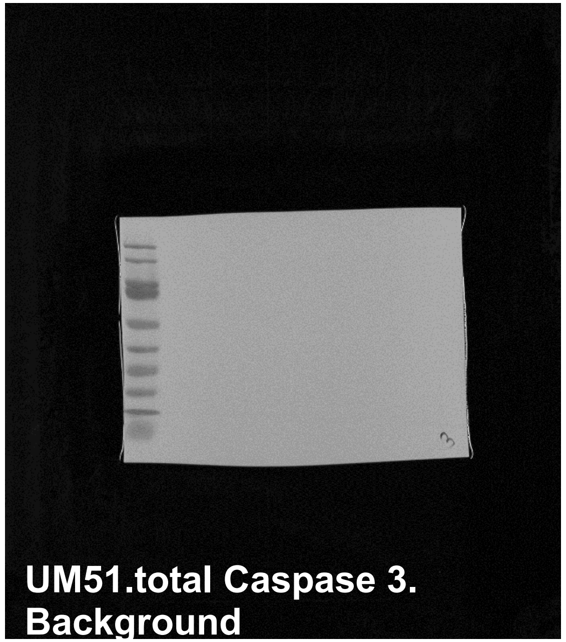


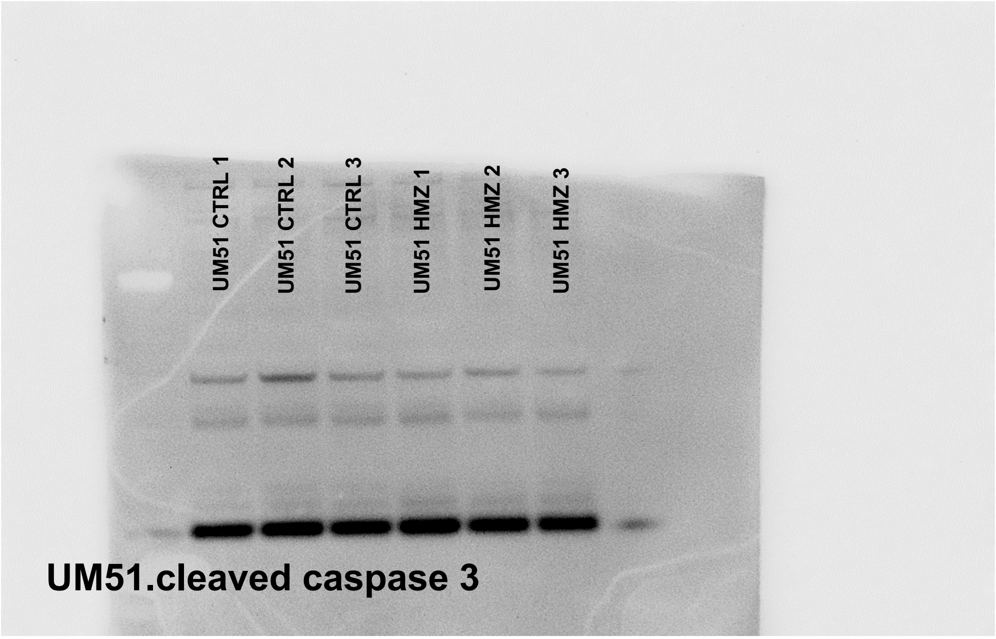


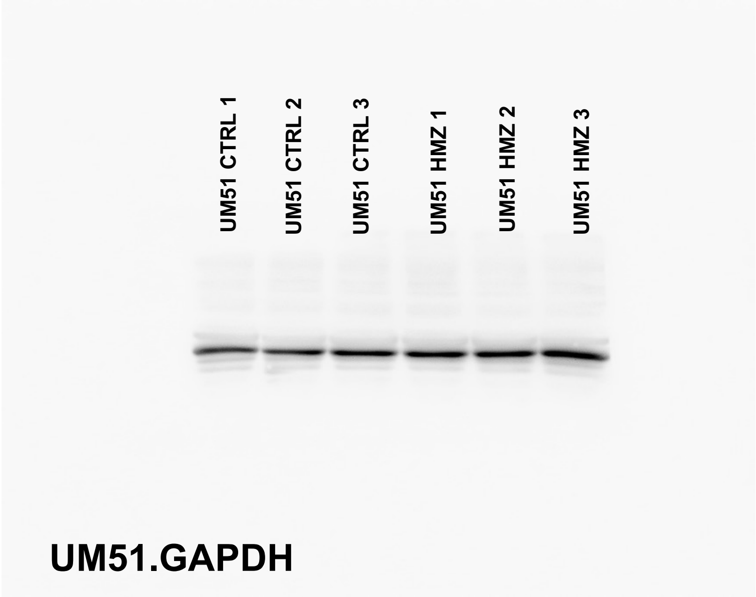


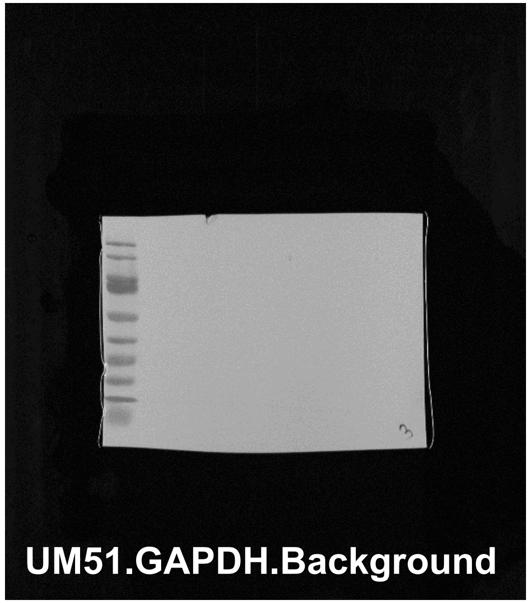


The whole blot membranes for supplementary figure 1. Cleaved caspase 3 was stained first, then the membrane was stripped and further used for total caspase 3.

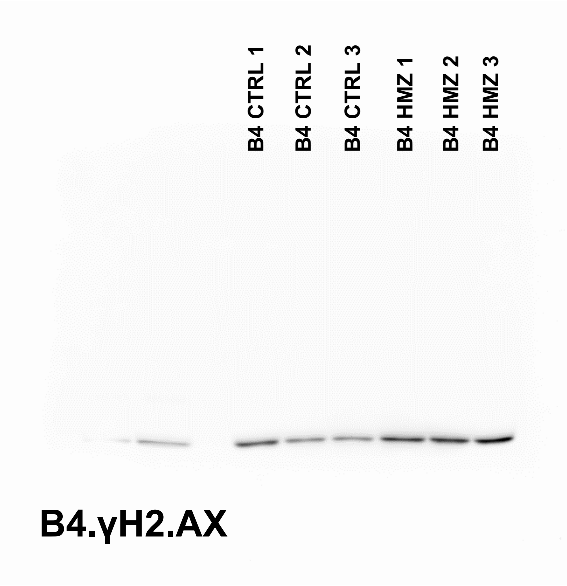


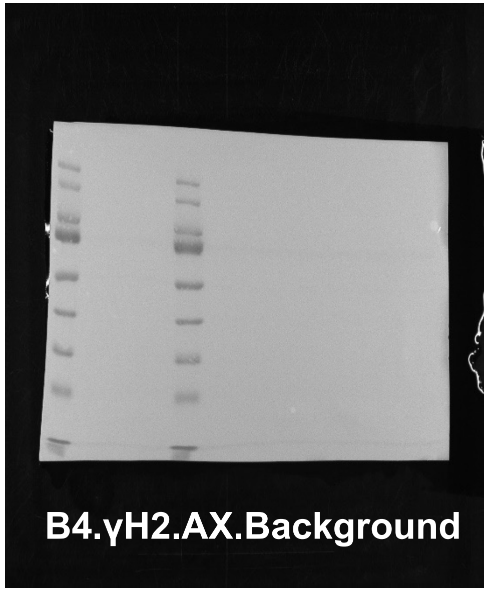


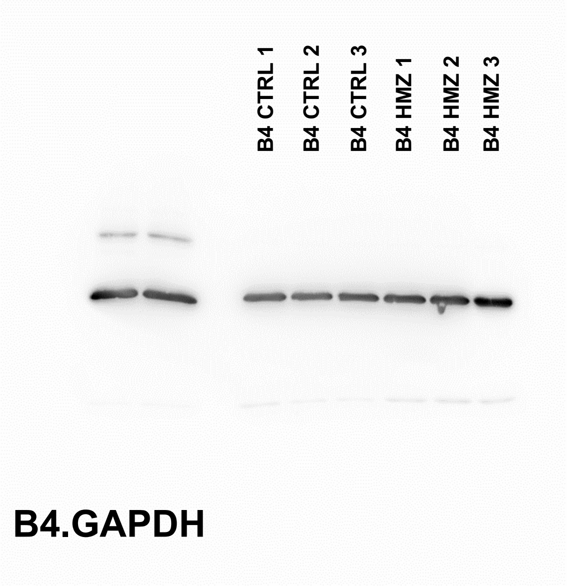


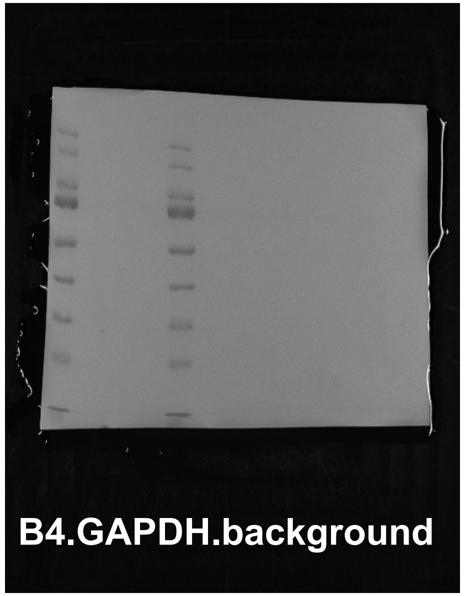


The whole blot membranes for main figure 3.

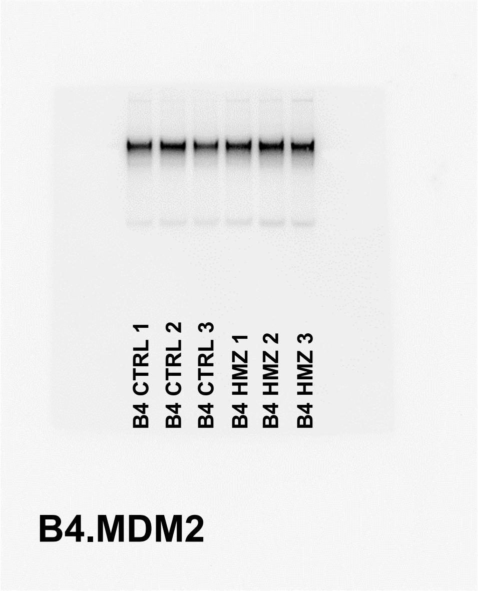


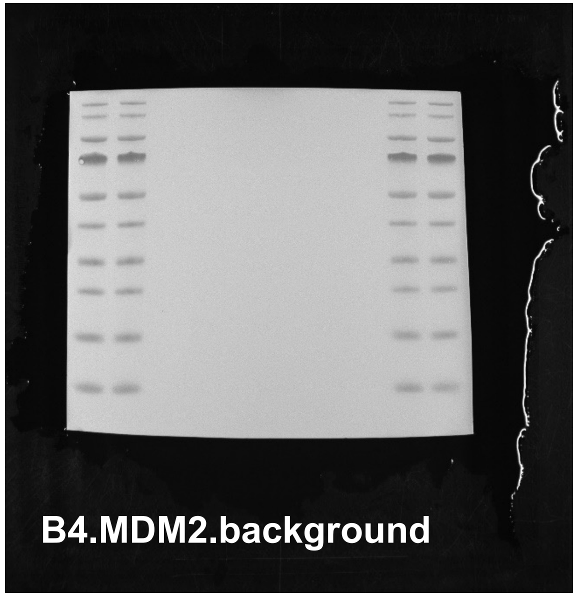


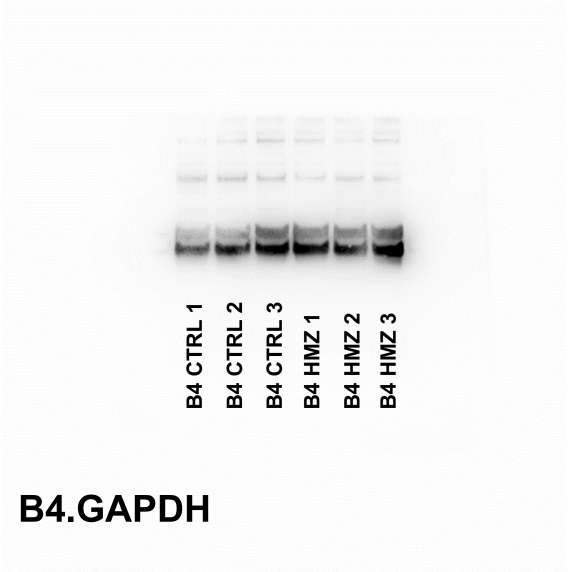


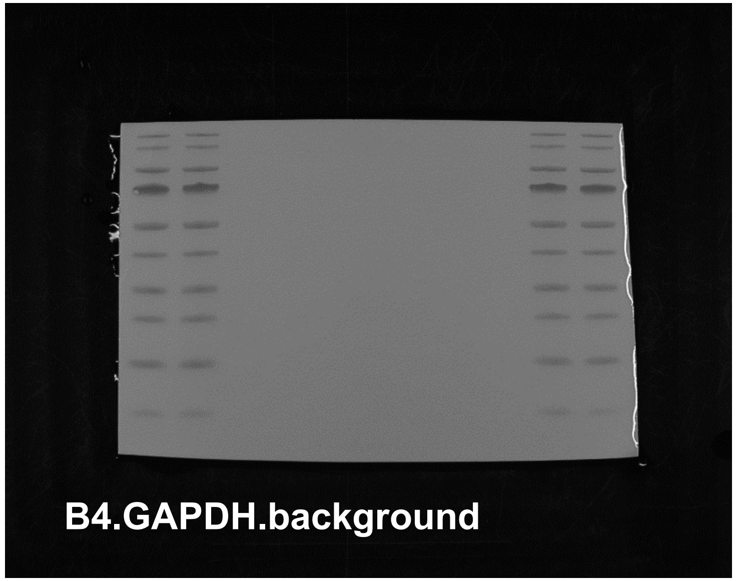


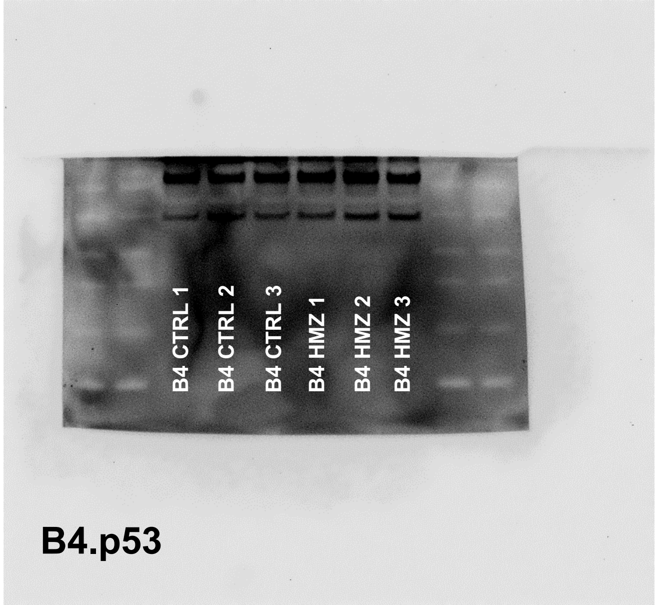


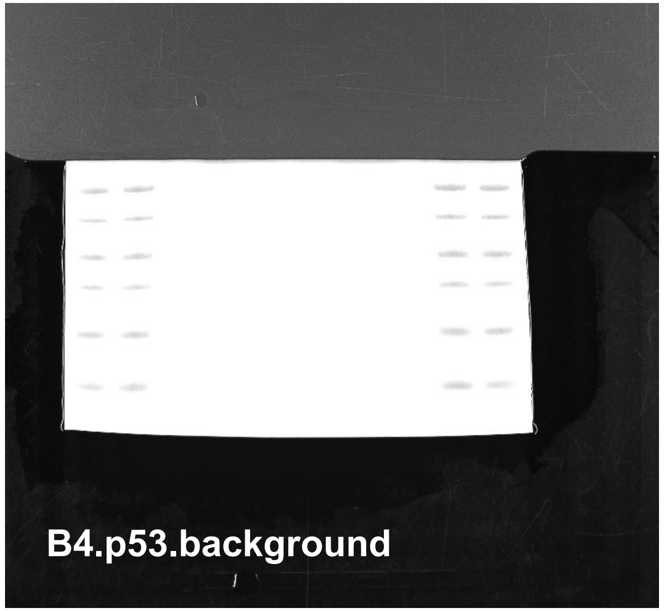


The whole blot membranes for main figure 4. The stainings were done on the same membrane for MDM2 and p53.

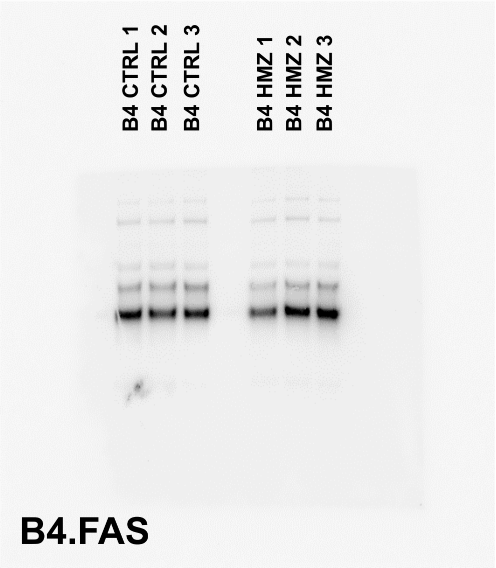


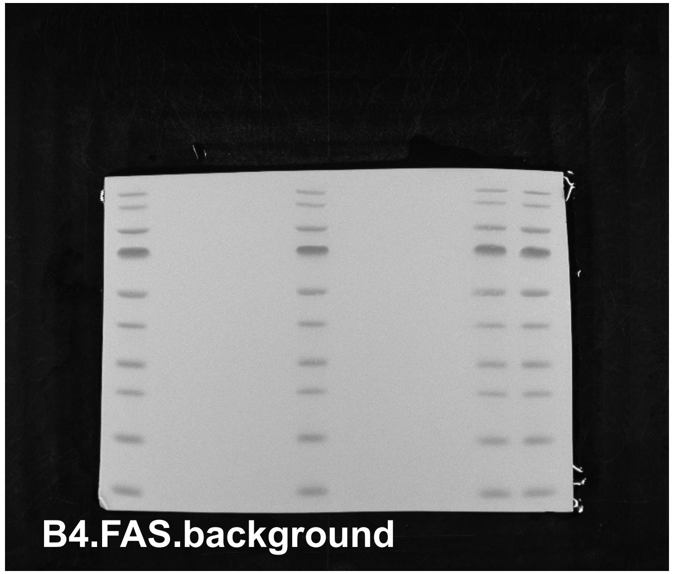


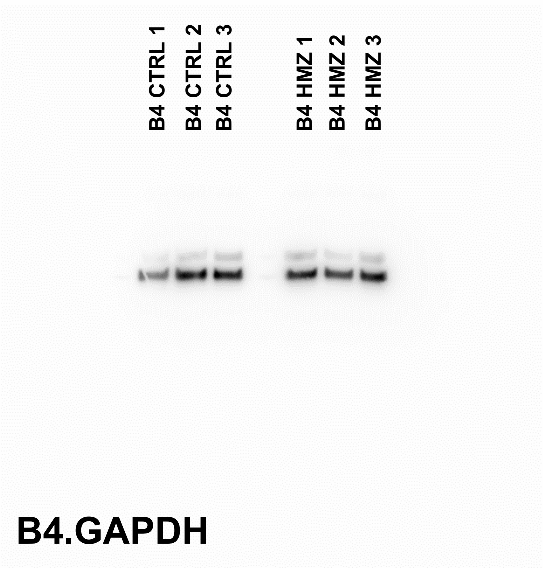


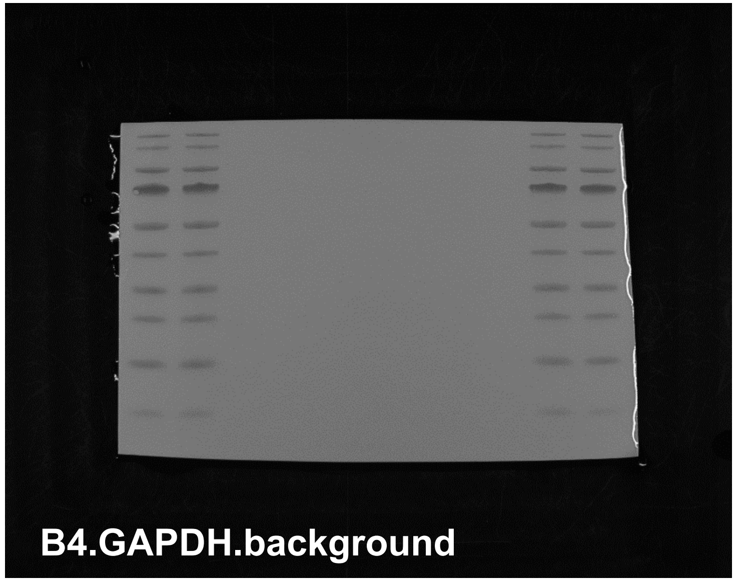


The whole blot membranes for main figure 4.

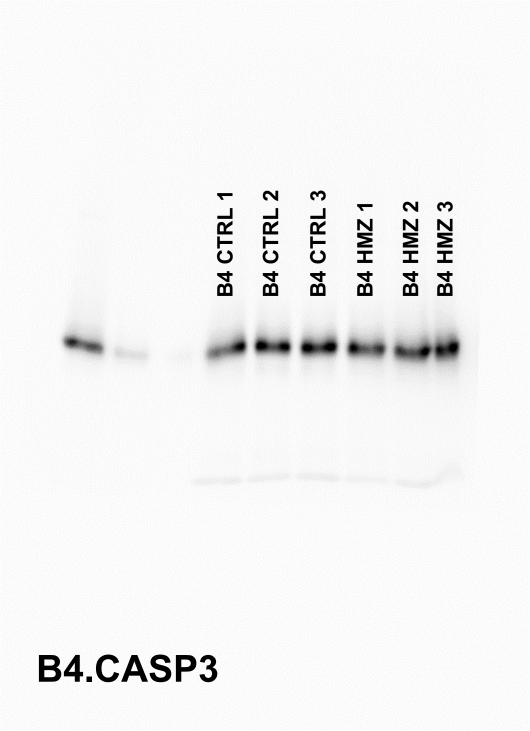


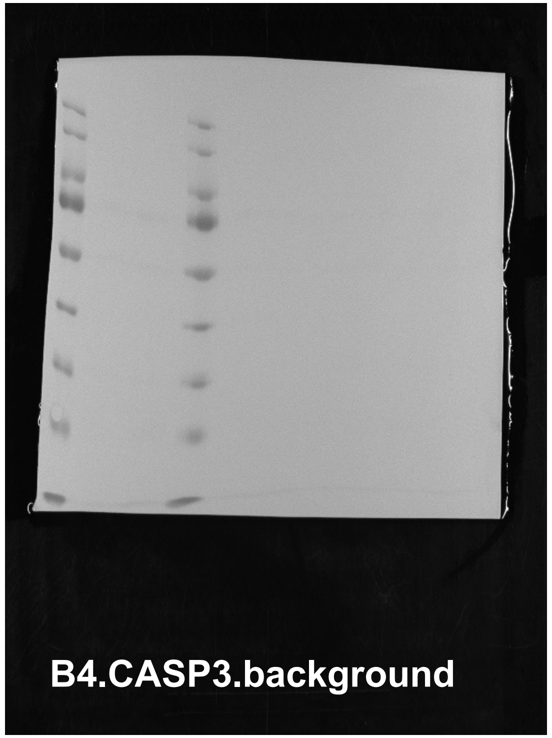


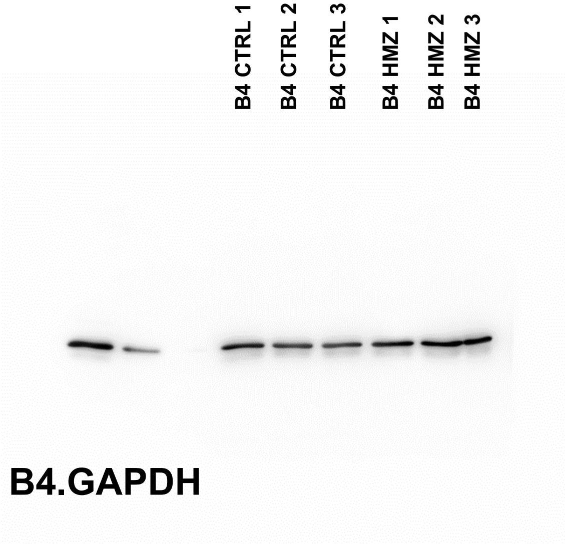


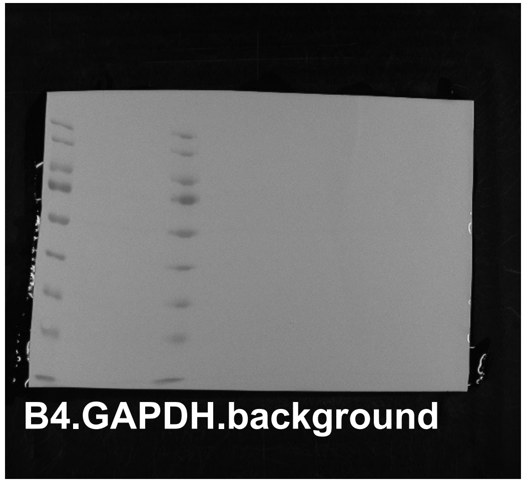


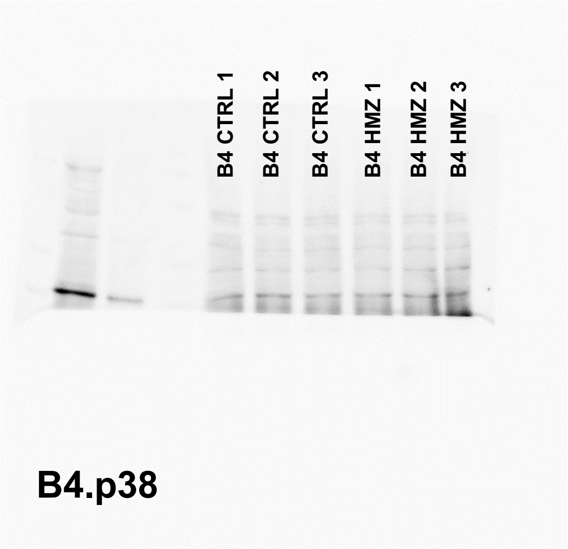


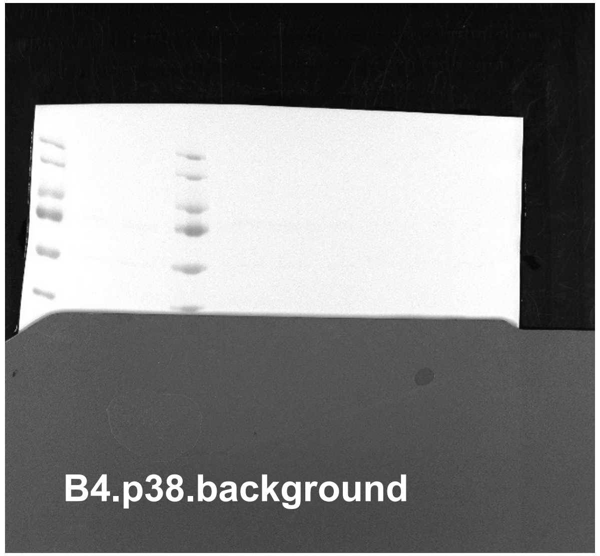


The whole blot membranes for main figure 4 and supplementary figure 1. The stainings were done on the same membrane for total p38 and CASP3.

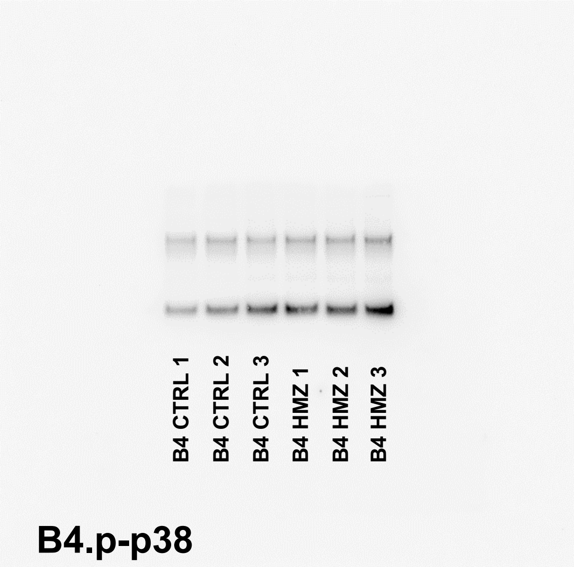


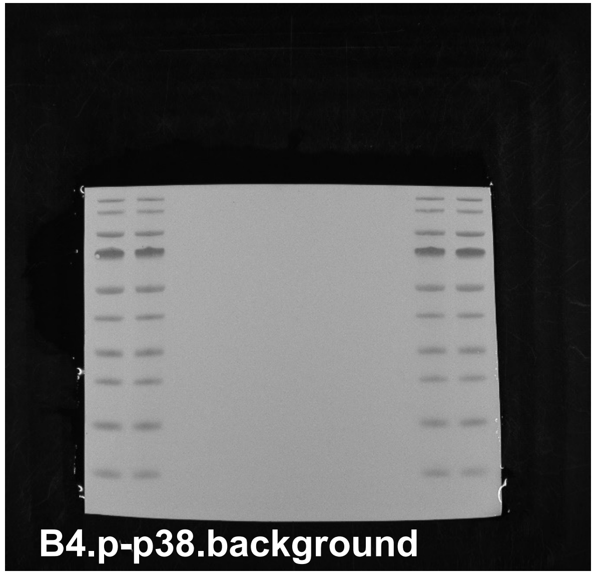


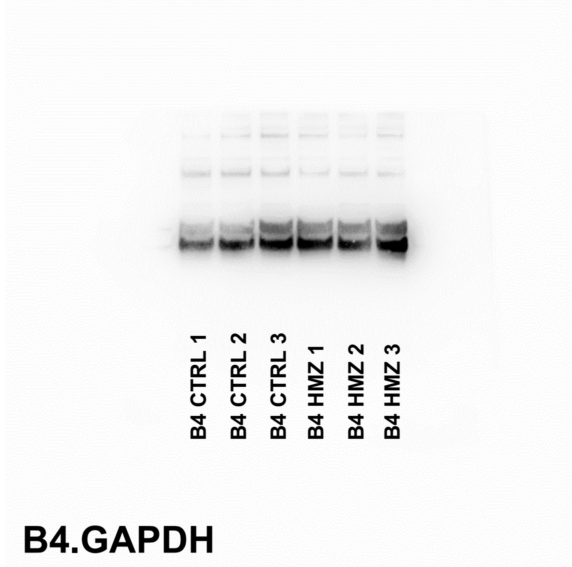


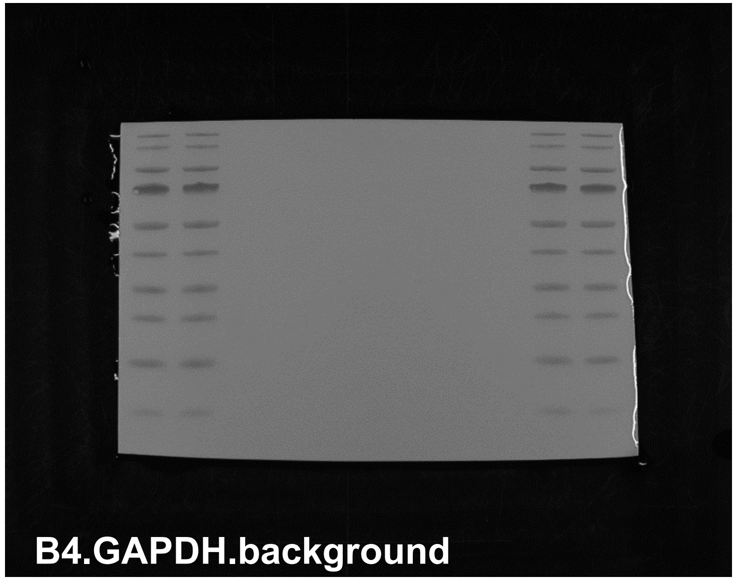


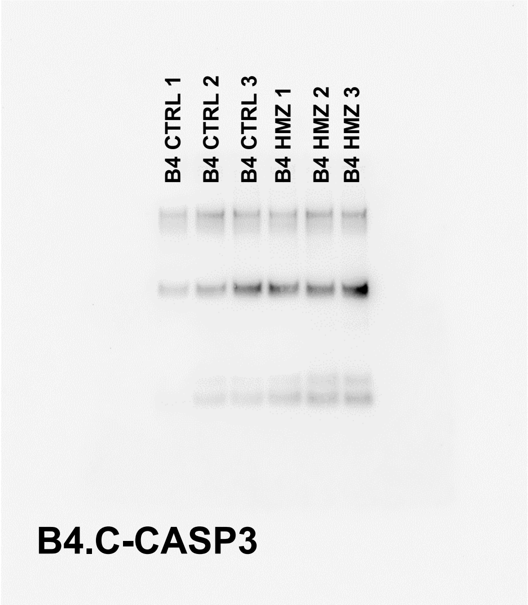


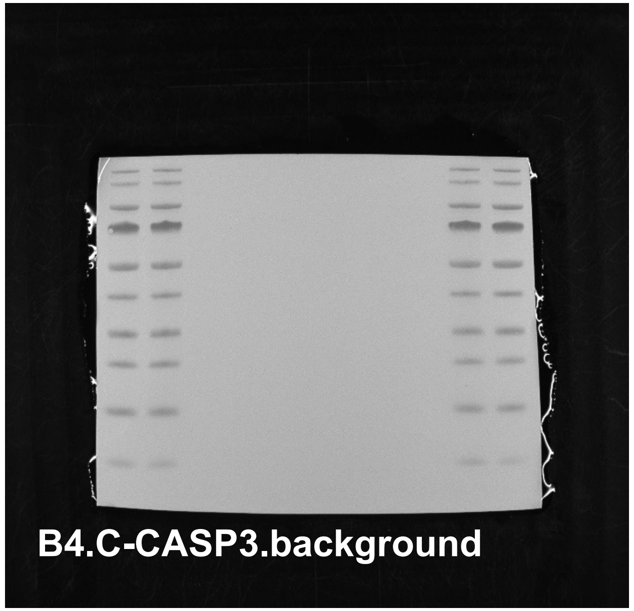


The whole blot membranes for main figure 4 and supplementary figure 1. The stainings were done on the same membrane for p-p38 and C-CASP3.

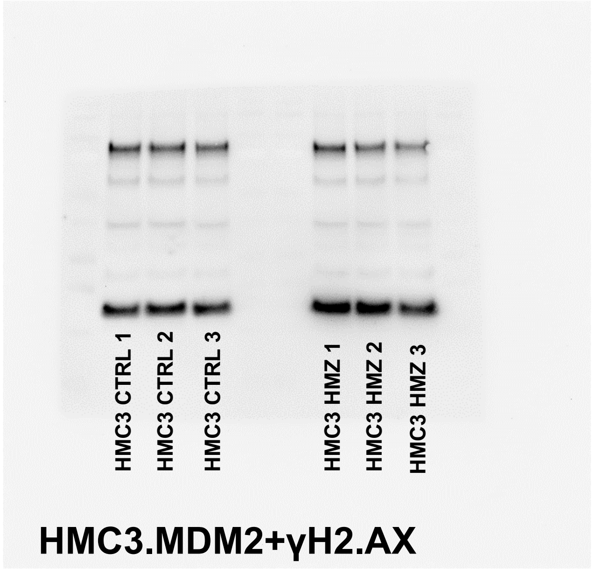


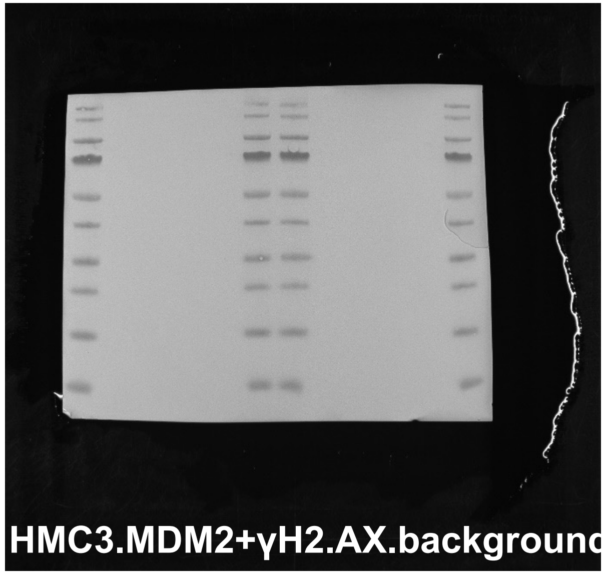


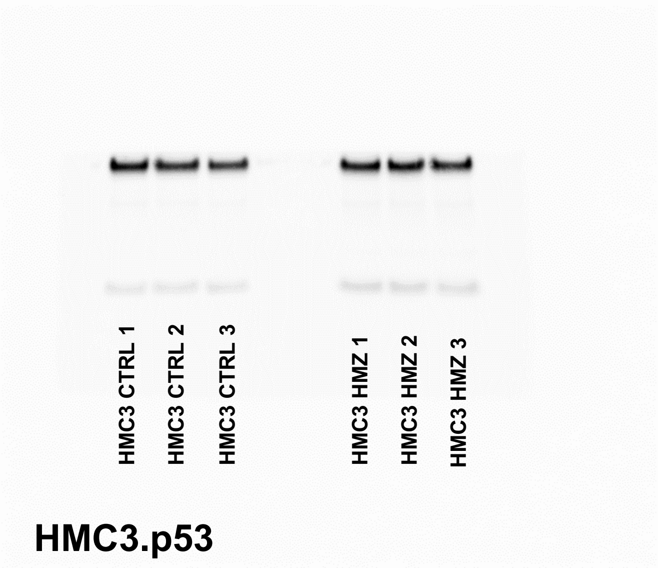


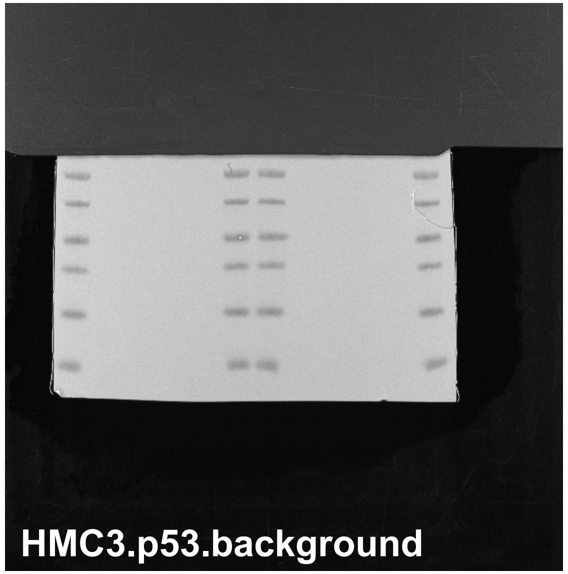


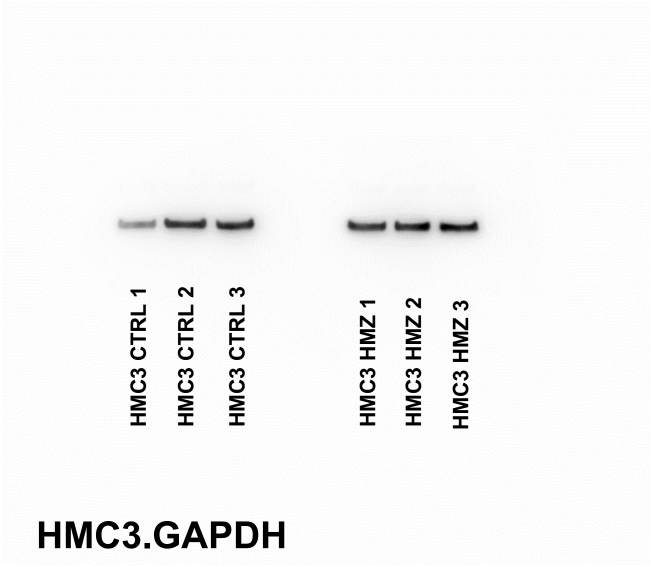


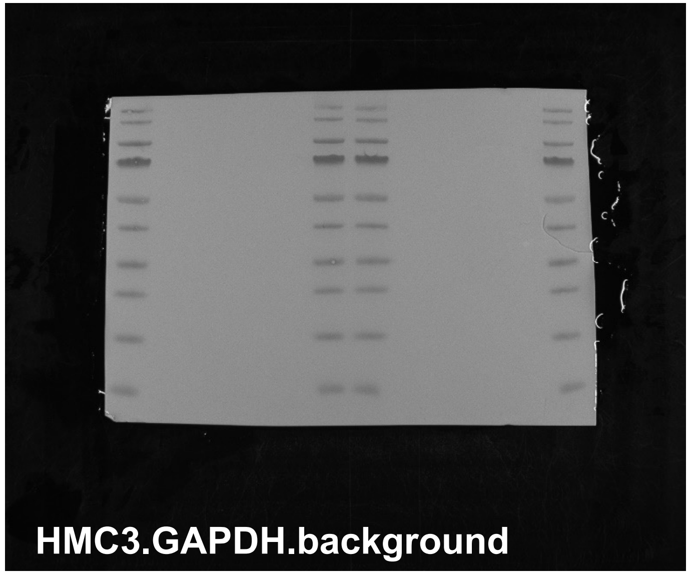


The whole blot membranes for supplementary figure 4. The stainings were done on the same membrane for all proteins.
